# DNA Methylation Patterns Expose Variations in Enhancer-Chromatin Modifications during Embryonic Stem Cell Differentiation

**DOI:** 10.1101/2020.11.25.397281

**Authors:** Adi Alajem, Hava Roth, Sofia Ratgauzer, Danny Bavli, Alex Motzik, Shlomtzion Lahav, Itay Peled, Oren Ram

## Abstract

In mammals, cellular identity is defined through strict regulation of chromatin modifications and DNA methylation that control gene expression. Methylation of cytosines at CpG sites in the genome is mainly associated with suppression; however, the reason for enhancer-specific methylation is not fully understood. We used sequential ChIP-bisulfite-sequencing for H3K4me1 and H3K27ac histone marks. By collecting data from the same genomic region, we identified enhancers differentially methylated between these two marks. We observed a global gain of CpG methylation primarily in H3K4me1-marked nucleosomes during mouse embryonic stem cell differentiation. This gain occurred largely in enhancer regions that regulate genes critical for differentiation. The higher levels of DNA methylation in H3K4me1-versus H3K27ac-marked enhancers, despite it being the same genomic region, indicates cellular heterogeneity of enhancer states. Analysis of single-cell RNA-seq profiles demonstrated that this heterogeneity correlates with gene expression during differentiation. Furthermore, heterogeneity of enhancer methylation correlates with transcription start site methylation. Our results provide insights into enhancer-based functional variation in complex biological systems.

**Author summary:** Cellular dynamics are underlined by numerous regulatory layers. The regulatory mechanism of interest in this work are enhancers. Enhancers are regulatory regions responsible, mainly, for increasing the possibility of transcription of a certain gene. Enhancers are marked by two distinct chemical groups-H3K4me1 and H3K27ac on the tail of histones. Histones are the proteins responsible for DNA packaging into condensed chromatin structure. In contrast, DNA methylation is a chemical modification often found on enhancers, and is traditionally associated with repression. A long debated question revolves around the functional relevance of DNA methylation in the context of enhancers. Here, we combined the two regulatory layers, histone marks and DNA methylation, to a single measurement that can highlight DNA methylation separately on each histone mark but at the same genomic region. When isolated with H3K4me1, enhancers showed higher levels of methylation compared to H3K27ac. As we measured the same genomic locations, we show that differences of DNA methylation between these marks can only be explained by cellular heterogeneity. We also demonstrated that these enhancers tend to play roles in stem cell differentiation and expression levels of the genes they control correlate with cell-to-cell variation.

## Introduction

Chromatin plasticity allows variation in transcriptional programs in cells with identical genetic codes. Cellular differentiation is regulated in part through modifications that decorate histone tails. Different modifications are associated with distinct genomic features such as active and repressive regions, distal elements, and promoters.

The histone marks regulate access of transcription factors to the genome, and hundreds of chromatin regulators read, write, and erase these marks. H3K4me1 and H3K27Ac are the main modifications of interest in this paper. H3K4me1 is situated on distal enhancer elements, whether they are poised or active. H3K27Ac predominantly marks active promoters and enhancers. When present together, the two modifications label active enhancers (1,2).

Chromatin immunoprecipitation followed by high-throughput sequencing (ChIP-seq) allows the profiling of different chromatin states using specific antibodies (3). To produce high-quality ChIP-seq maps large numbers of cells are required, and the output map represents an average signal over the bulk population masking cellular heterogeneity. Recently, we and others have developed single-cell ChIP-seq methods (4–6); however, antibody efficacy and barcoding limitations in most cases prohibit in-depth single-cell analyses (4,6,7). Imaging of single nucleosomes is possible (8), but there is currently no robust method that combines single nucleosome imaging with sequence analysis. Therefore, the ability to directly link between chromatin and DNA methylation states is limited.

Methylation of cytosines (C) at the 5th position to form 5-methylcytosine (5mC) in the context of CpG dinucleotides controls lineage-specific expression of developmental genes (9), genomic imprinting (10), X chromosome inactivation (11), and retrotransposon silencing (12). The 5mC modification is deposited and maintained by DNA methyltransferases (DNMTs) DNMT1, DNMT3A, and DNMT3B, and an instrumental part in 5mC removal are the dioxygenases Tet1, Tet2, and Tet3 (13,14). Loss of any component of this regulatory system results in global changes in DNA methylation. Although embryonic stem cells (ESCs) retain their ability to self-renew after the loss of any of the proteins of the methylation machinery (15), loss of any one protein does impair the ability of ESCs to properly differentiate in culture (16–18), and loss of all three Tet proteins causes embryonic lethality (19–21).

Mapping cytosine methylation levels on a genome-wide scale is usually done by bisulfite sequencing (BS-seq). Bisulfite treatment of DNA converts unmethylated cytosines to uracils, which are mapped to a reference genome to distinguish 5mCs from Cs at a single-nucleotide resolution (22). It is important to note that bisulfite conversion cannot detect 5hmC from 5mc. However, 5hmc levels are extremely low (<0.05%) in ESCs (19) and therefore negligible in our analyses.

Most of the CpGs in the genome are located outside regulatory regions. Active promoters are mostly unmethylated regions (UMRs), whereas *cis*-regulatory sequences such as enhancers show methylation levels ranging from 20% to 80% and are referred to as low-methylated regions (LMRs) (23). Characterization of methylation levels is mostly evaluated by analysis of bulk populations. This poses the possibility that LMRs reflect heterogeneity of methylation in enhancer regions. The development of Single cell DNA methylation profiling (24–26) could answer the question, however, these technologies suffer from low sensitivity and false negative measurements (27). Particularly, regions with few CpG dinucleotides such as enhancers are not detected (28). Therefore, this hypothesis is difficult to test. On top of that, the relationship between histone modifications and DNA methylation is unclear, leaving the role of DNA methylation in specific chromatin contexts an open question.

To better read this complex code, it is critical to approach the subject with high resolution and take cellular heterogeneity into account. Here we used sequential ChIP-bisulfite-sequencing (ChIP-BS-seq) to directly measure CpG methylation under a specific chromatin modification. We analyzed pluripotent ESCs and differentiated cells with H3K4me1 and H3K27ac antibodies to detect chromatin regions associated with enhancers. Transcribed genes were identified using H3K36me3 antibodies, and poised promoters and enhancers were detected using previously published H3K27me3 data (29). We found that enhancers gain methylation during differentiation. Distinctively, DNA methylation accumulates to a greater extent within H3K4me1-marked nucleosomal DNA than within H3K27ac-marked nucleosomal DNA. Surprisingly, when the same enhancer is immunoprecipitated with H3K4me1, DNA methylation levels are higher than when it is immunoprecipitated with H3K27Ac, despite it being the same genomic region. We call these differentially methylated enhancers (DMEs). **These DMEs reflect cellular heterogeneity of enhancer states, and, as shown using single-cell RNA-seq, this heterogeneity is reflected at the mRNA level during ESC differentiation**. The DME heterogeneity also correlates with transcription start site (TSS) methylation, providing a possible explanation for changes in TSS methylation states during differentiation. The method described in this paper directly links between chromatin states and DNA methylation levels. Sequential ChIP bisulfite followed by high throughput sequencing complements single cell methods by overcoming sensitivity limitations, particularly highlighting heterogeneous enhancer states.

## Results

### Direct measurement of DNA methylation in enhancers

To study enhancers in the context of DNA methylation, we cultured mouse ESCs for 2 days in medium lacking inhibitors of MEK and GSK3 (‘2i’), previously shown to down-regulate global DNA methylation without compromising pluripotency (30). In order to compare pluripotent enhancers to enhancers of a differentiated state, mouse ESCs were differentiated using 4-day treatment with retinoic acid (31). This differentiation protocol promotes a broad spectrum of cellular differentiation states, producing epiblast stem cell-like (EpiSC) states and extra embryonic endoderm (XEN) cells (32) (Fig. S1). Analysis of ESCs and differentiated cells was performed using ChIP-BS sequencing. Genome-wide sequencing of DNA treated by bisulfite is a standard approach for mapping 5mC in an unbiased manner. However, as many cells are pooled together prior to BS-seq, the resultant maps average 5mC levels, blind to differences between cells and to variation of chromatin states in a given genomic location. In order to overcome this limitation, we profiled 5mc levels at enhancers after precipitating the DNA with anti-H3K4me1 and anti-H3K27ac antibodies (33) (Fig. 1A).

**Figure 1.**
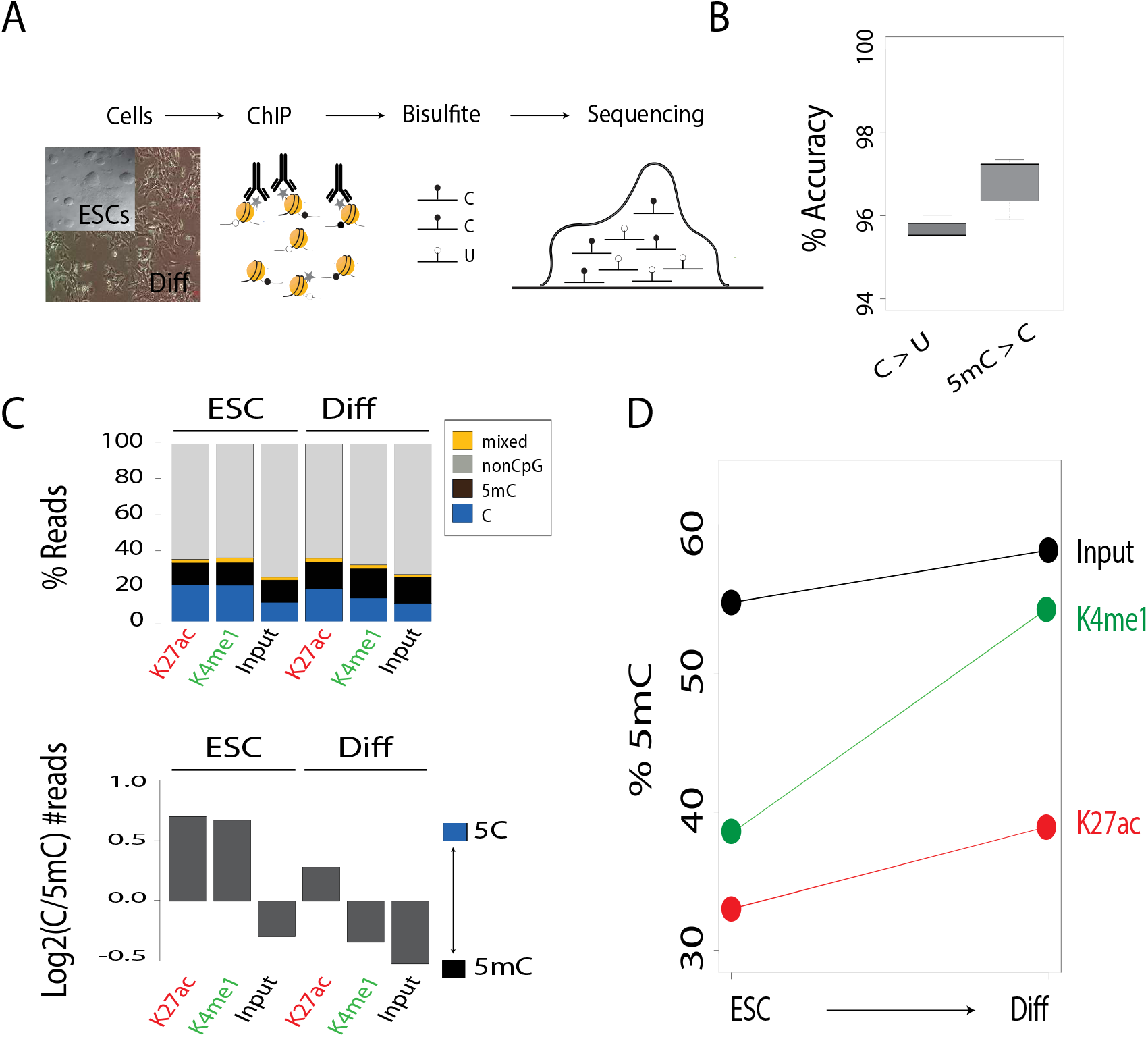
DNA methylation increases during differentiation. **(A)** Schematic representation of the ChIP-BS-seq-based analysis of mouse ESCs and differentiated cells (Diff). **(B)** Quality of BS-seq was analyzed by spiking in oligonucleotides with 5mC and 5C. Plotted is accuracy (%) for each nucleotide. **(C)** Upper panel: Bar plot of percentages of C, 5mC, mixed (in which adjacent CpGs had different methylation states), and non-CpG reads for ChIP experiments and input for ESCs and differentiated cells. Lower panel: Log2 fold change between C and 5mC reads for each condition. **(D)** Percentage of 5mC in ChIP experiments and input in ESCs and differentiated cells.

To control for bisulfite efficacy, PCR and sequencing errors, we repeated each experiment twice and each sample was spiked with a synthetic oligonucleotide containing both methylated and unmethylated cytosines. Measurements of the spiked-in oligonucleotide showed an accuracy of more than 95% for both unmethylated and methylated cytosines (Fig. 1B). ChIP-BS-seq maps showed that enhancer regions are largely depleted of CpG dinucleotides (34). Nevertheless, it is possible for two or more CpG dinucleotides to be present on the same read. Be it the case and DNA methylation is detected, we expect all CpGs on the same read to be methylated due to non-specificity of DNMTs in such small a scale. Should only one be methylated, we removed the read from further analysis (4% of the reads). We considered it a result of bisulfite inefficiency/PCR/sequencing error (Fig. 1C, top, mixed).

As we wanted to compare our findings to results achieved using bulk ChIP-BS, we added a whole cell genomic DNA extract from both ESCs and differentiated samples (marked as input) to each experiment. Genomic regions positive for H3K4me1 or H3K27ac in ESCs showed lower 5mC levels than the same regions in the input sample (Fig. 1C, bottom). This observation aligns with the hypothesis of heterogeneity in chromatin states at a given loci. Both H3K4me1 and H3K27ac mark active chromatin regions, whereas the input sample is an unbiased genome-wide measurement that includes active, repressed and unmarked regions.

Methylation levels in differentiated cells were higher in all three chromatin contexts (H3K4me1, H3K27ac, and the input sample) than in ESCs. Moreover, in H3K4me1 samples, both differentiated cells and ESCs showed higher levels of 5mc per read (each read was counted as either methylated or unmethylated) than in H3K27ac (Fig. 1C, bottom). Similar trends were observed for single 5mC percentages (Fig. 1D). This suggests that during differentiation H3K4me1 is more 5mC permissive than H3K27ac.

### LMRs reflect variation in chromatin states of enhancers

To maximize our ability to profile CpG methylation of precipitated regions, we segmented the mouse genome into 200-bp bins, the length of the nucleosomal DNA unit. We analyzed bins that included reads covering at least 30 unique CpGs counts (Table S1 and S2), which were located at least 10 kb away from TSSs. To compare changes in methylation states upon differentiation, we extracted 111,377 H3K4me1 and 16,019 H3K27ac genomic bins overlapping with enhancer peaks as analyzed by the HOMER peak algorithm (35). Notably, at distal enhancer regions an increase in 5mC levels upon differentiation was observed for both H3K27ac and H3K4me1. This rise was much more profound for H3K4me1 than for H3K27ac (Fig. 2A and 2B, student’s t. test p<e^−10^).

**Figure 2.**
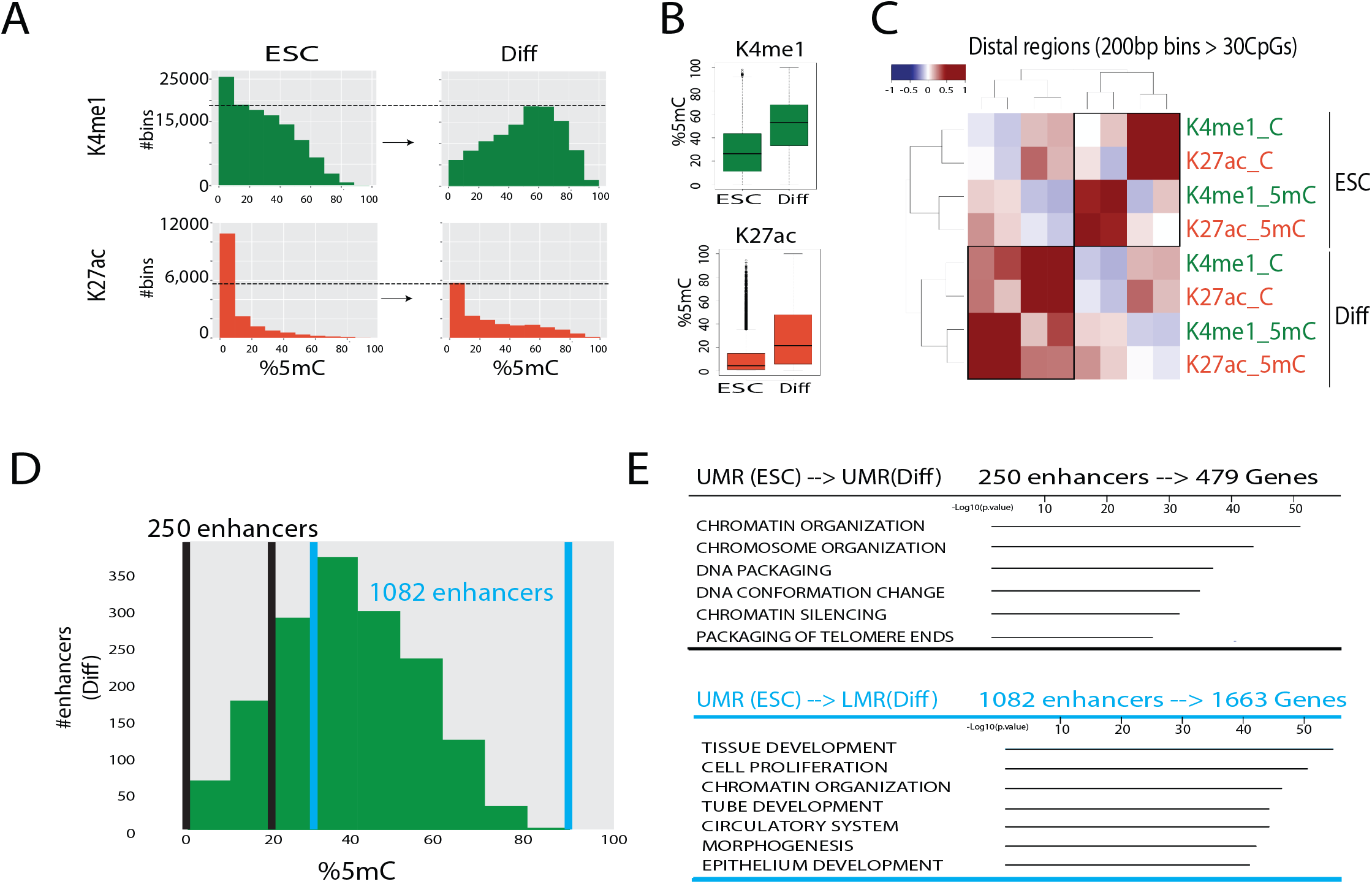
Enhancer-specific DNA methylation occurs during differentiation. **(A)** Histograms of number of bins with indicated 5mC percentages for H3K4me1 and H3K27ac in ESCs and differentiated cells (Diff) over distal enhancers; percentages were calculated in 200-bp genomic segments. **(B)** Boxplots of 5mC percentages in H3K4me1-bound regions (n=111,377) and H3K27ac-bound regions (n=16,019) in ESCs and differentiated cells. Thick lines indicate the medians. The boxes range from 25%-75%, and the whiskers extend to ±1.5 of the interquartile range. **(C)** Spearman rank correlations between H3K4me1 and H3K27ac ChIP-seq maps of distal enhancers for ESCs and differentiated cells, separating signals of C vs. 5mC levels (#reads) within 200-bp genomic bins. **(D)** Histogram of 5mC% in regions bound by H3K4me1 in differentiated cells that were UMRs in ESCs; percentages were calculated in 200-bp bins. **(E)** MsigDB Gene Ontology terms (66) of genes regulated by enhancers that remained in the UMR state after differentiation (upper table) and of enhancers that were UMRs in ESCs and became LMRs in differentiated cells (lower table). Only top enriched GO categories were shown. Complete gene list can be found in Supplementary table S3.

Enhancer regions precipitated with H3K4me1 or with H3K27ac have lower levels of CpG methylation than the same genomic locations evaluated by input BS-seq (Fig. S2). This was expected as input samples include a mixture of nucleosomal states (active, primed, poised, and silenced). In contrast, the 5mC signal within transcribed genes, evaluated by ChIP-BS-seq using H3K36me3, was higher than the detected 5mC signal in the input BS-seq samples (36) (Fig. S2). This aligns with previous reports showing association of CpG methylation with transcribed regions (37).

To further explore enhancer-specific methylation during differentiation, we divided each of the four ChIP BS-seq maps obtained (H3K4me1 and H3K27ac, from ESC and differentiated cells) into two groups: unmethylated (C) and methylated (5mC). To compare between DNA methylation levels in each chromatin context and cell line, we performed a Pearson correlation test between all eight groups. In ESCs, a strong correlation was found between H3K4me1_C and H3K27ac_C, as well as between H3K4me1_5mC and H3K27ac_5mC. However, weak to no correlation was found between methylated and unmethylated reads regardless of modification (Fig. 2C). This result emphasizes that in ESCs, enhancer central peaks are largely unmethylated, hence the strong correlation. The CpG methylated reads come from enhancer flanking regions which are different between the two marks. In differentiated cells, a higher correlation was observed between unmethylated and methylated states (Fig. 2C). This suggests that upon differentiation, CpG methylation moves into enhancer central regions. This aligns with our hypothesis that upon differentiation the same enhancers can be found in two states, active (unmethylated) and repressed (methylated) as indicated by LMR states. For a given enhancer, a subset of cells can be in an unmethylated and active state (H3K4me1 and H3K27ac) while in another subset of cells the same enhancer can be methylated, probably in the process of inactivation (H3K4me1 only).

Enhancers are associated with LMRs as previous genome-wide bisulfite profiling indicates (23). However, the functional relevance of this intermediate methylation state is not known. The length of LMRs ranges between 400bp to 2 kb (Fig. S3). Data from our enhancer-specific ChIP coupled to bisulfite sequencing suggests that regions of apparently low methylation reflect mixing of subpopulations. In our Sequential-BS-seq, some enhancers were identified as LMRs (>20% 5mC). Despite that, a substantial number of enhancers were unmethylated. In ESCs, approximately 45% of enhancers contained a UMR. Upon differentiation, only about 15% of enhancers had UMRs, supporting our hypothesis of mixed enhancer states. We also found that overlap between H3K4me1 and H3K27ac UMRs is more common in ESCs than in differentiated cells.

To further test whether LMRs result from a mixture of subpopulations, we collected all H3K4me1 enhancers which predominantly were UMRs in ESCs (<20% 5mC). We then divided them into two groups: H3K4me1 enhancers that maintained an unmethylated state upon differentiation (250 enhancers) and those that shifted their methylation state from UMR in ESCs to LMR upon differentiation (1,082 enhancers with 5mC >30%) (Fig. 2D). Next, we linked the enhancers with their target genes using the mouse *cis*-regulatory atlas (38). Enhancers that remain unmethylated upon differentiation tend to be associated with genes involved in cell maintenance, chromatin, and chromosome packaging. In contrast, enhancers that became LMRs upon differentiation are associated with genes involved in differentiation including *Notch*, *Wnt*, and *Gata* (Fig. 2E). This strongly suggests that LMR enhancers reflect cell-to-cell variation during differentiation.

### Distinct methylation of enhancers identifies chromatin-based subpopulations

To further analyze enhancer states and possible cellular subpopulations, we analyzed ChIP-BS-seq data to compare H3K4me1 to H3K27ac enhancer-specific nucleosomes (Fig. 3A). If the same histone modifications were homogenously present in all cells over a particular enhancer region, we expected to see no differences in DNA methylation levels regardless of the antibody used for the ChIP assay. DNA from the same genomic region but with different 5mC levels that immunoprecipitated with antibodies of different histone marks can only be explained by the presence of subpopulations of cells with different chromatin states.

**Figure 3.**
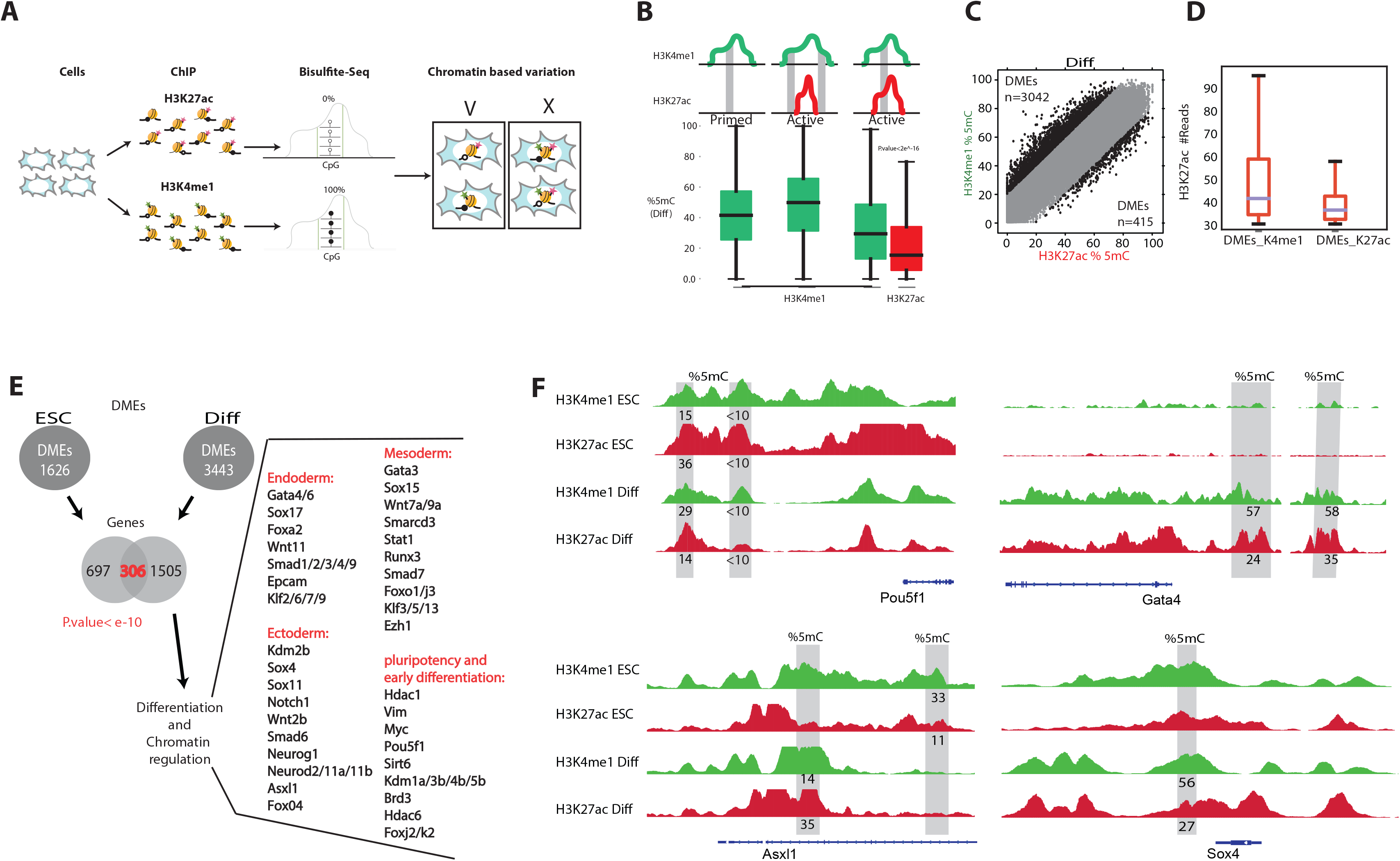
DNA methylation states of distal enhancers support chromatin-based cellular heterogeneity. **(A)** Schematic representation of ChIP-BS-seq for H3K4me1 and H3K27ac histone marks emphasizing that different 5mC levels are explained only by chromatin-based heterogeneity. **(B)** Locations of 5mC in the three most frequent scenarios (schematics above the boxplots). Boxplots show 5mC percentages in each chromatin state. **(C)** Plot of 5mC percentages in H3K4me1 and H3K27ac genomic segments in differentiated cells. Only genomic bins with more than 30 CpGs in both ChIP-BS-seq assays are shown. DMEs were defined as 200-bp bins with methylation differences exceeding 20%. **(D)** H3K27ac read counts within DMEs bound by H3K4me1 (left) and H3K27ac (right). **(E)** Genes with DMEs annotated based on the mouse *cis*-regulatory atlas (38). The Venn diagram shows a significant overlap of 306 genes (Hypergeometric test, p<e^−10^) of genes regulated by DMEs in both undifferentiated and differentiated cells. Some of the overlapping genes are listed, grouped according to function. **(F)** IGV tracks of DMEs for *Pou5f1*, *Gata4*, *Asxl1*, and *Sox4*. Percentages of 5mC in the grey blocks are specified beneath each track. For adjacent DMEs, 5mC levels were averaged.

In ESCs, we detected 356,366 H3K4me1-enriched bins and 66,210 H3K27ac-enriched bins; 55,537 of them overlapped (Table S1). In differentiated cells, 257,891 H3K4me1-enriched bins and 64,080 H3K27ac-enriched bins were detected, and 57,093 of them overlapped. (Table S2). In ESCs, overlapping bins included about 50% of total enhancers, and in differentiated cells the overlapping bins covered 60% of total enhancers. The enhancers not represented were mostly poorly enriched in the ChIP assay, or had too low CpG densities for our assay to detect.

In differentiated cells, enhancer bins that passed minimal coverage of 5mC mapped most frequently to three types of histone-bound regions. In one group, only H3K4me1 bins were found. In another group, both H3K4me1 and H3K27ac show ChIPseq signals, but the detected bins came from H3K4me1 distinct regions. In the third group, the most relevant to our downstream analysis, both H3K4me1 and H3K27ac signals exist and 5mC bins from the same location were extracted for both histone marks (Fig. 3B). In ESCs, 5mC levels in regions marked by both modifications showed a slight but significant difference in abundances of H3K4me1 and H3K27ac (student’s t. test p<e^−10^) (Fig. S4). However, upon differentiation, in the same genomic regions, 5mC levels were strikingly higher when the regions were bound by H3K4me1-marked nucleosome than H3K27ac nucleosomes (Fig. 3B, student’s t. test p<e^−10^). Again, this finding is a strong indication of chromatin-based cellular heterogeneity. In addition, H3K4me1 covered a higher fraction of the genome than H3K27ac, and regions containing only H3K4me1 peaks showed higher methylation levels than those containing both marks. This suggests that H3K4me1-only enhancers might be more susceptible to deactivation by methylation.

To identify enhancer regions where methylation levels correspond to a unique chromatin state, we needed to account for errors in bisulfite modification, PCR, and sequencing. For bisulfite conversion, we know that the error rate is 2-5% (based on synthetic spike-in, Fig. 1B). Therefore, to ensure that a region is accurately defined as differentially methylated, we decided to extract bins with at least 20% difference in 5mC levels between the two histone marks (Fig. 3C). There are two types of DMEs. Those in which 5mC levels are higher in H3K4me1 and those in which 5mC levels are higher in H3K27ac. In ESCs there was a similar number of DMEs of both types: H3K4me1 DMEs (n=946) and H3K27ac DMEs (n=680) (Fig. S5). This symmetry, together with overall low levels of 5mC, suggests that DMEs do not explain the variation of cellular states in ESCs. Upon differentiation, we noticed that most of DMEs tend to be more methylated when immunoprecipitated with H3K4me1 as opposed to H3K27ac. Overall, 3,042 DMEs had higher methylation levels when pulled-down with H3K4me1, while only 415 DMEs had higher methylation levels when pulled-down with H3K27Ac. Moreover, H3K4me1 DMEs showed a strong tendency to be associated with H3K27ac peak centers. In cases in which DMEs showed higher methylation levels in H3K27ac than H3K4me1, they were more likely to be found outside peak centers (Fig. 3D, student’s t. test p<e^−10^). As central peaks better associate with TF binding regions (39), we speculate that enhancer deactivation is manifested primarily through removal of H3K27ac, which can then be traced by DNA methylation accumulation under H3K4me1.

To map enhancers to their target genes we used the previously reported mouse *cis*-regulatory atlas (38). Overall, 1,003 genes in ESCs and 1,811 genes in differentiated cells were regulated by enhancers containing DMEs (Fig. 3E and Table S4). These genes were significantly enriched for cell cycle, chromatin, and differentiation of the three main germ layers (Fig. 3E). For example, Gata4/6 (40,41) are master regulators of XEN cells. Sox4 (42), Asxl1 (42) and Smad (43) genes are regulators of the EpiSC and act as "pioneer" factors that open the compacted chromatin for other proteins at certain enhancer sites. The enhancers of these genes are differentially methylated and expression of the genes was activated only upon differentiation (Fig. 3F). This suggests that differentiation using retinoic acid promotes differentiation to EpiSC and XEN programs in distinct subpopulations of cells.

Cellular heterogeneity reflected by DMEs might arise from allelic asymmetry (28). To rule out the possibility that DMEs reflect allelic asymmetry, we analyzed DMEs in the X chromosome compared to autosomes. The R1 ESC line is of male origin; therefore, if allelic asymmetry indeed underlies our observations of differential methylation in enhancer regions, the number of DMEs on the X chromosome should be significantly lower or non-detected as a result of the fact that these cells contain only one X chromosome. However, we found no evidence for deviation in DME distributions between autosomes and the X chromosome (Fig. S6), suggesting that allelic asymmetry is not a major cause of DMEs.

### DMEs regulate genes associated with cell-to-cell variation

To determine whether DMEs are indeed linked to genes that drive cell-to-cell variation during differentiation, we used the InDrop method (44). We collected single-cell RNA-seq data from 869 ESCs and 696 differentiated cells, and detected about 10,000 transcripts per cell. We identified approximately 2,000 genes in ESCs and approximately 3,000 genes in differentiated cells (Fig. S7). ESCs expressed similar levels of *Oct4* (Fig. 4A), emphasizing their pluripotent states. Using the Seurat algorithm (45), in ESCs, we detected a spectrum of cellular states associated with Nanog expression levels (Fig. 4A). In more detail, they had different levels of *Nanog* and other genes such as Tulp3 and mitochondrial genes emphasizing their tendency towards a ground state (46) (Fig. S8). Cells that express little or no *Nanog* and in this case higher levels of *Set*, *Gnl3*, *Hmgb2*, and *Hsp90ab1* may be in a more primed cellular state (47,48).

**Figure 4.**
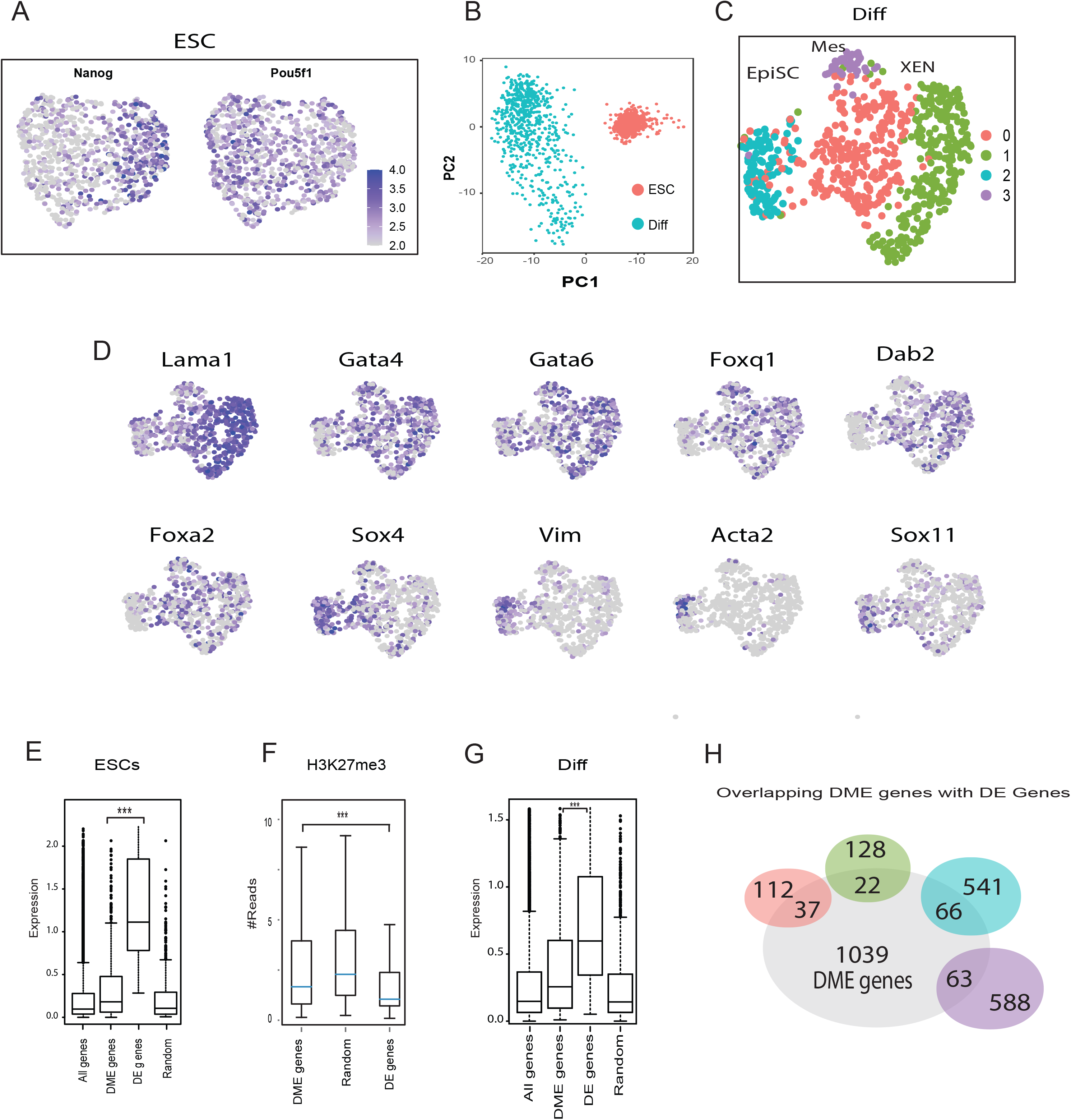
Single-cell RNA-seq indicates that DMEs regulate cellular heterogeneity. **(A)** Single-cell RNA-seq data from 869 ESCs was processed using the InDrop method. UMAP plots and unbiased *k*-means clustering revealed two subpopulations explained, mostly, by *Nanog* expression levels while Pou5f1 expression levels remained stable in most cells. Levels vary from low expression in gray to high expression in purple. **(B)** Single-cell RNA-seq data of ESCs and differentiated cells was subjected to principal component analysis. Distances between cells in the plot indicate similarities between transcriptional profiles. **(C)** Single-cell RNA-seq data from 696 differentiated cells was processed using the InDrop method. UMAP and unbiased *k*-means clustering revealed four differentiated subpopulations separated by color. Each dot represents an individual cell. Clusters 0 and 3 show upregulation of genes associated with mesendodermal progenitors. Cluster 1 has high levels of XEN cells markers. Cluster 2 shows EpiSC markers. **(D)** Expression levels of 10 representative genes shown by UMAPs of differentiated cells with low expression in gray to high expression in purple **. (E)** Boxplot of expression levels in ESCs of all genes, genes regulated by DMEs, differentially expressed genes (DE genes) revealed by single cell RNA-seq analysis and a random set of genes. DME genes and DE genes differ in expression levels (student’s t. test p<e^−10^). **(F)** H3K27me3 reads around TSS regions (1 kb on either side of the TSS) in ESCs, showing that genes regulated by DMEs and randomly selected genes are more repressed compared to DE genes (student’s t. test p<e^−10^). **(G)** Boxplot of expression levels in differentiated cells of all genes, genes regulated by DMEs and DE genes and a random set of genes. A significant difference in expression levels between DME genes and DE genes was found (student’s t. test p<e^−10^). **(H)** Illustration of the intersection between genes regulated by 1227 DMEs, and the four clusters of genes identified using single cell RNA-seq analysis. DMEs are shown in gray and single cell genes are colored according to their clusters (panel E) (Hypergeometric tests, p<e^−10^).

For differentiated cells, single-cell RNA-seq results were indicative of a higher degree of cellular heterogeneity than observed in ESCs (Fig. 4B). In differentiated cells, four clusters with different upregulated markers were identified (Fig. 4C and table S5). Representative gene markers are shown in Figure 4D. Clusters 0 and 3 show upregulation of genes associated with mesendodermal progenitors such as Foxa2 (53). Cluster 1 has high levels of *Lama1, Gata4*, *Gata6*, *Foxq1 and Dab2* which are markers of XEN cells (40,49,50). Cluster 2 represents EpiSC with markers such as *Vim*, *Sox4*, *Sox11* and *Acta2* (51,52).

Single-cell RNA-seq mostly detects highly expressed genes, whereas chromatin profiling enables detection of repressed, poised, and active genes. In ESCs, most of the genes regulated by DMEs are expressed at low levels (Fig. 4E). These genes are likely poised for activation due to promoter states known as bivalent domains that contain both H3K27me3 and H3K4me3 (53). Indeed, promoters of the genes predicted to be regulated by DMEs showed significantly higher H3K27me3 levels compared to randomly selected and differentially expressed genes (Fig. 4F). In differentiated cells, genes regulated by DMEs are expressed at higher levels than randomly selected genes (Fig. 4G, student’s t. test p<e^−10^). Moreover, DMEs of genes that drive differentiation have significant overlap with marker genes of the four subpopulations identified by single-cell RNA-seq. Overall, 188 of the 1369 genes detected by the single-cell RNA-seq assay were regulated by DMEs (Fig. 4H, hypergeometric test p<e^−10^). This analysis emphasizes the power of DME profiling to complement RNA-seq analyses which tends to capture only highly expressed genes.

### TSS proximal regions of genes regulated by DMEs are highly methylated during differentiation

In pluripotent ESCs, genes that regulate differentiation are transcriptionally poised through deposition of H3K27me3 over regulatory regions such as TSSs and enhancers (53,54). Our data indicates that H3K27ac marks active enhancers and TSSs (Fig. 5A and 5B). H3K4me1 nucleosomes do not peak around TSSs but rather mark TSS proximal regions, which in many cases possess enhancers that regulate transcription from the adjacent TSS (1,54). To evaluate DNA methylation in TSS proximal regions, we divided each chromatin map into unmethylated versus methylated CpG reads. H3K27ac nucleosomes peaked in the 1.5-kb region around TSSs, and these regions were found to be depleted of 5mC (Fig. 5A). In contrast, H3K4me1 nucleosomes bind diffusely about 5 kb around TSS regions. In undifferentiated cells, DNA bound by H3K4me1-marked nucleosomes was overall unmethylated (Fig. 5A); however, upon differentiation, 5mC accumulated predominantly on H3K4me1 proximal regions (Fig. 5A). In addition, H3K4me1-5mC patterns positively correlate (Pearson correlation test, r=0.82) with expression as measured using single-cell RNA-seq (Fig. 5B and C and Fig. S9A and S9B).

**Figure 5.**
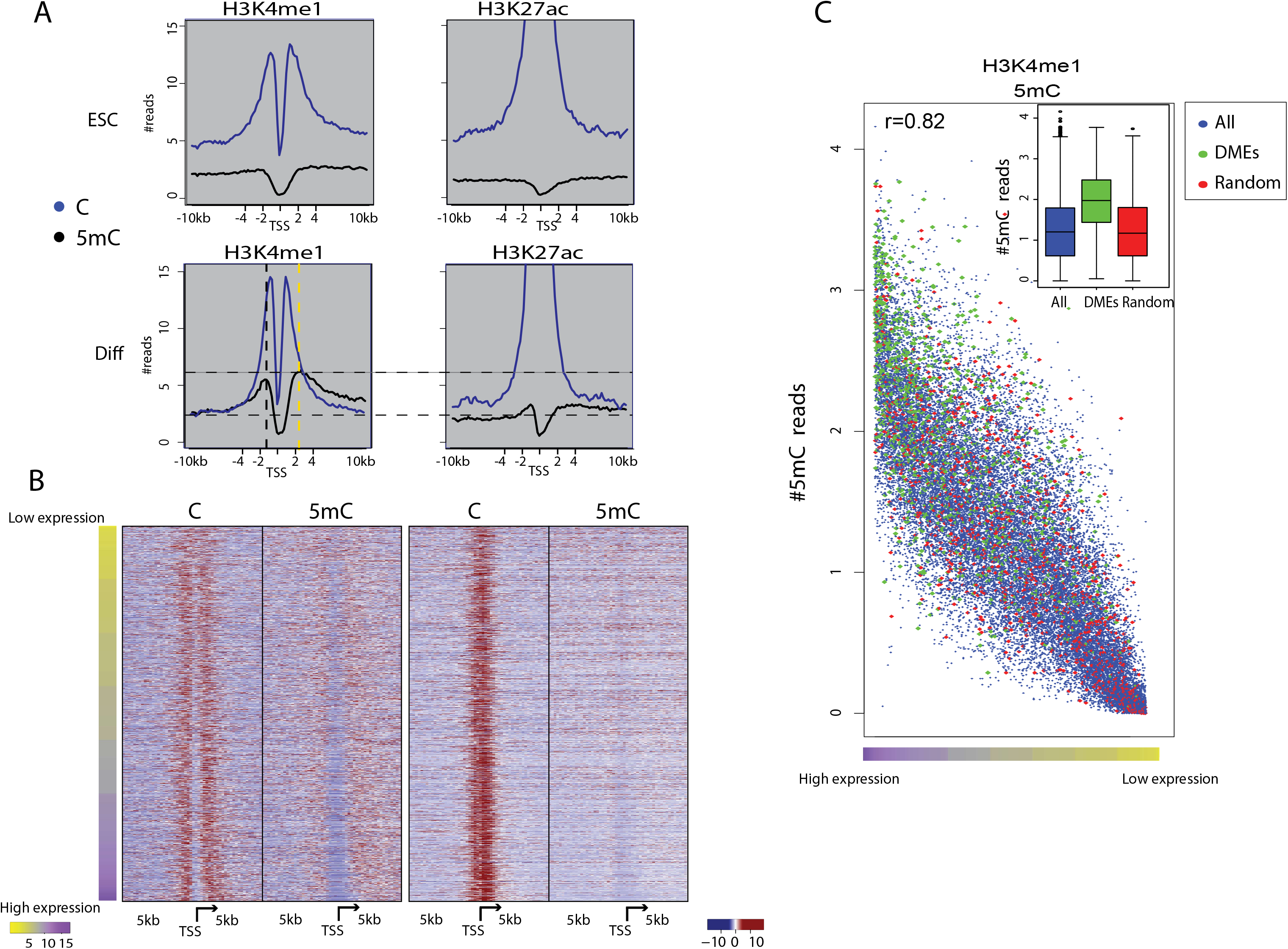
DMEs are associated with high H3K4me1-5mC levels in regions proximal to transcriptional start sites. (**A**) Composite plots of unmethylated (blue) vs. methylated (black) reads of H3K27ac and H3K4me1 maps over ±10 kb around the TSSs of ESCs and differentiated cells. Vertical dashed lines represent 5mC peak centers (upstream to TSS in black and downstream to TSS in yellow). Horizontal dashed lines represent minimum and maximum of 5mC peaks. (**B**) Heatmaps showing normalized numbers of 5C and 5mC aligned reads within 200-bp bins in the region 5 kb up and downstream of TSSs. To reduce the number of TSSs presented, we randomly selected ~3,500 representative TSSs from ten different expression bins as measured by single-cell RNA-seq. (**C**) Normalized number of 5mC reads around TSSs precipitated with anti-H3K4me1 from differentiated cells plotted from high expression to low expression (left to right on the x axis) for all genes (black), DME-regulated genes (green), and a randomly selected set of genes equal in size to the DME set (red). Boxplots summarizing the same data are shown in the upper right side.

If DMEs reflect cellular variation due to differentiation, we hypothesized that we would find higher levels of DNA methylation also in proximal regions of genes that are regulated by DMEs. Indeed, TSS proximal regions regulated by DMEs had dramatically higher levels of methylation compared to other TSS proximal regions as well as to a randomly selected set of regions (Fig. 5C, student’s t. test p<e^−10^). This methylation elevation was observed both upstream and downstream of the TSS, ruling out the possibility that methylation is simply a reflection of higher methylation in actively expressed regions. We found a regulatory connection between distal DMEs and proximal enhancer regulation, hence, strengthening the notion that DNA methylation on H3K4me1 is associated with cellular variation.

## Discussion

Abnormalities in DNA methylation at enhancers are associated with developmental defects and disease (55–57). How DNA methylation at enhancers influences the poised, primed, active, or silenced state of genes is not completely understood (2). Methylation of cytosine in the context of CpG dinucleotides is associated with enhancer repression (58,59); however, analyses of enhancer states of bulk cells average heterogeneity and, therefore, cannot directly correlate methylation state with gene expression. In order to achieve this level of depth in analysis, it is required to combine between single-cell ChIP-seq and whole genome bisulfite sequencing. Single-cell technologies suggest that heterogeneity across cells is manifested specifically through changes within regulatory regions such as enhancers (4,60). However, due to sensitivity limitations of single-cell assays, they cannot illuminate whether LMR states measured in bulk mix active and repressed enhancer states, or reflect a permissiveness of DNA methylation within active regions.

Dynamic changes in enhancer methylation depend on DNMTs, cell proliferation, and transcription factor binding. Two mechanisms of action have been suggested for DNMTs. These enzymes may be actively involved in the removal of transcription factors from enhancers or may passively act on sites free from transcription factors to ensure enhancer decommissioning (23,61). Previous genome-wide measurements showed that active chromatin regions are associated with very little to no CpG methylation (62–64). Therefore, the assumption was that these DNA regions are physically protected from DNMTs due to binding of other factors. However, we show that many enhancers of actively transcribed genes are within the LMR range. This observation strengthens the assertion of cellular heterogeneity.

To determine the reason behind LMRs observed in enhancers, it was essential to extract enhancer regions of known regulatory states. As active enhancers are mostly associated with H3K4me1 and H3K27ac histone marks, we measured DNA methylation in regions associated with each of these two histone marks by combining sequential chromatin immunoprecipitation with bisulfite sequencing. Surprisingly, when H3K4me1 and H3K27Ac are measured for a given enhancer, we found that LMRs are mostly associated with H3K4me1 and not with H3K27Ac. This suggests that LMRs reflect the heterogeneity of different enhancer states between cells.

Enhancers of genes associated with linage differentiation are UMRs for both marks in ESCs and LMRs upon differentiation. These LMRs are found to be enriched with DMEs, which are significantly more methylated when pulled-down with H3K4me1 antibodies than with H3K27ac antibodies. This suggests a temporal regulatory scheme: usually, during cell differentiation, active enhancers are marked with both H3K4me1 and H3K27ac. We show that in a subset of cells, some enhancers are decommissioned, H3K27ac is reduced and the DNA is left bound by H3K4me1-marked nucleosomes, acquiring CpG methylation in the process. Our observation that there are higher levels of H3K4me1 methylation in regions depleted of H3K27ac supports this hypothesis, however, additional experiments are required to prove that DNA methylation induces H3K27ac removal.

To determine the effects of DMEs on gene expression, we applied single-cell RNA-seq analysis. In ESCs, differentiating genes were largely repressed through H3K27me3, which explains the overall deficiency of DMEs. However, upon differentiation, DNA methylation becomes a dominant factor in enhancer regulation. In differentiated cells genes regulated by DMEs overlap with genes identified as differentially expressed in single-cell RNA-seq. This suggests that once differentiation is initiated, DMEs become functional.

Sequential ChIP-bisulfite data reduces noise originating from cells that lack a given enhancer or that are diverse in their enhancer chromatin state (e.g., an enhancer that in some cells is marked only by H3K4me1 but in other cells is marked by both H3K4me1 and H3K27ac). In addition, this method allowed quantification of methylation in low-density CpG regions that cannot be detected using the reduced representation bisulfite sequencing method (65). The Sequential ChIP-bisulfite method described here allowed us to dissect how epigenomic information influences gene expression, improving our ability to accurately map enhancer states. It also highlighted chromatin variation that cannot be captured using common ChIP-seq. Sequential ChIP-bisulfite can be used to identify dynamic biological processes with respect to enhancer regulation. Combined with single-cell RNA-seq, it enables the analysis of enhancer states in the context of gene expression. Our study provides a path towards a better understanding of dynamic enhancer regulation, which could be beneficial for the study of dysfunctional gene regulation that occurs during cancer development and aging.

## Acknowledgments

O.R. is supported by research grants from the European Research Council (ERC, # 715260 SC-EpiCode), the Israeli Center of Research Excellence (I-CORE) program, the Israel Science Foundation (ISF, #1618/16), and the Azriely Foundation Scholar Program for Distinguished Junior Faculty. The authors extend special thanks to Prof. Eran Mesorer, Dr. Yosi Buganim, and Dr. Maayan Salton from the Hebrew University for helpful discussions and critical reading of the manuscript and to Dena Herrmann Ennis for manuscript proofreading.

## Author Contributions

O.R., S.R. and A.A. wrote the manuscript. O.R. S.L. and A.A conceived the study and prepared the figures. O.R. and A.A. designed the experiments. A.A. and S.R. performed ChIP-BS-seq, tissue culturing, differentiation, and library preparation for ChIP-BS-seq. D.B., A.M. and A.A. prepared the microfluidcs and performed single cell RNA-seq experiments. O.R., H.R., and I.P. performed the computational analyses of ChIP-BS-seq and single-cell RNA-seq data.

## Author Information

All ChIP-BS-seq and single-cell RNA-seq data have been deposited in the Gene Expression Omnibus database (GEO) under accession number (GSE158378). For reviewers: please use this link: https://www.ncbi.nlm.nih.gov/geo/query/acc.cgi?acc=GSE158378. The authors declare no competing financial interests. Correspondence and requests for materials should be addressed to O.R.

## Material & Methods

### Cells

R1 ESCs were grown on 0.1% gelatin-coated dishes and maintained in DMEM with 15% ESC-grade fetal calf serum, 50 μg/mL penicillin, 50 μg/mL streptomycin, 2 mM L-glutamine, 1 mM sodium pyruvate, 0.1 mM nonessential amino acids, 0.1 mM β-mercaptoethanol, and 1000 U/ml LIF, supplemented with 1 μM PD0325901, an inhibitor of the MEK/ERK pathway, and 3 μM CHIR99021, an inhibitor of GSK3. For retinoic acid-induced differentiation, cells were grown on 0.1% gelatin-coated dishes for 4 days in DMEM with 10% ESC-grade fetal calf serum, 50 μg/mL penicillin, 50 μg/mL streptomycin, 2 mM L-glutamine, 1 mM sodium pyruvate, 0.1 mM nonessential amino acids, 0.1 mM β-mercaptoethanol, and 1 μM retinoic acid.

### Micrococcal nuclease digestion

Cells were trypsinized, washed with PBS, and resuspended in 100 μL PBS per 1×10^6^ cells. The same volume of lysis buffer (100 mM Tris-HCl [pH 8], 300 mM NaCl, 2% Triton X-100, 0.2% sodium deoxycholate, 10 mM CaCl2, and EDTA-free protease inhibitor cocktail added at a ratio of 1:100 [Sigma]) with 50 U/mL MNase (Thermo Scientific) was added, and the cells were incubated for 10 min on ice and then for 15 min at 37 °C. Reactions were stopped by addition of 20 mM EGTA followed by centrifugation (20,000 g, 2 min). DNA was purified from the supernatants using Ampure XP beads (Beckman Coulter) and electrophoresed (Fig. S10).

### ChIP

Antibodies (H3K4me1 abcam ab8895, H3K27Ac abcam ab4729, H3K36me abcam ab9050, 2 μg antibody per 2×10^7^ cells) were incubated with 25 μL protein A Dynabeads (Thermo Scientific) previously washed twice with blocking buffer (0.5% BSA, 0.5% Tween-20 in PBS) for 2 h at 4 °C in blocking buffer. The conjugated beads were washed twice with blocking buffer before adding the lysate. Lysates were prepared as follows: Cells were trypsinized, washed with PBS, and resuspended with 500 μL PBS per 2×10^7^ cells. The same volume of lysis buffer with 100 U/mL MNase was added, and the cells were incubated for 10 min on ice and then for 15 min at 37 °C. Reactions were stopped by adding 20 mM EGTA followed by centrifugation (20,000 g, 2 min). Supernatants were added to the conjugated beads, and samples were incubated overnight with rotation. Supernatants were removed, and the beads were washed twice with RIPA buffer (10 mM Tris-HCl [pH 8], 140 mM NaCl, 1% Triton X-100, 0.1% sodium deoxycholate, 0.1% SDS, 1 mM EDTA), twice with RIPA buffer high salt (10 mM Tris-HCl [pH 8], 360 mM NaCl, 1% Triton X-100, 0.1% sodium deoxycholate, 0.1% SDS, 1 mM EDTA), twice with LiCl wash buffer (10 mM Tris-HCl [pH 8], 250 mM LiCl, 0.5% sodium deoxycholate, 1 mM EDTA, 0.5% IGEPAL CA-630 [Sigma]), and twice with 10 mM Tris-HCl [pH 8]. The beads were resuspended in 45 μL 10 mM Tris-HCl [pH 8] and 20 μg RNase A. The beads were incubated for 30 min at 37 °C, and 60 μg proteinase K was added. The beads were further incubated for 2 h at 37 °C. Proteinase K was inactivated by heating to 65 °C for 15 min, and 50 μL elution buffer (10 mM Tris-HCl [pH 8], 300 mM NaCl, 1% Triton X-100, 0.1% sodium deoxycholate, 0.1% SDS, 1 mM EDTA) was added, and the samples were incubated in 65 °C for 15 min. DNA was purified from supernatants using Ampure XP beads (Beckman Coulter). DNA was eluted with 22 μL 10 mM Tris-HCl [pH 8].

### ChIP-BS library preparation

To create spiked DNA sequences that contain both 5mC and C, we used oligonucleotide primers H1 and H2 and the Spike-in template (Table 1). Primers H1 and H2, which contain unmodified C, were used to amplify Spike-in using d(5mCTP) rather than dCTP in the dNTP mix to produce an amplicon that contains 5mC in the central region and C in the primer regions. DNA was purified using Ampure XP beads.

**Table 1.**
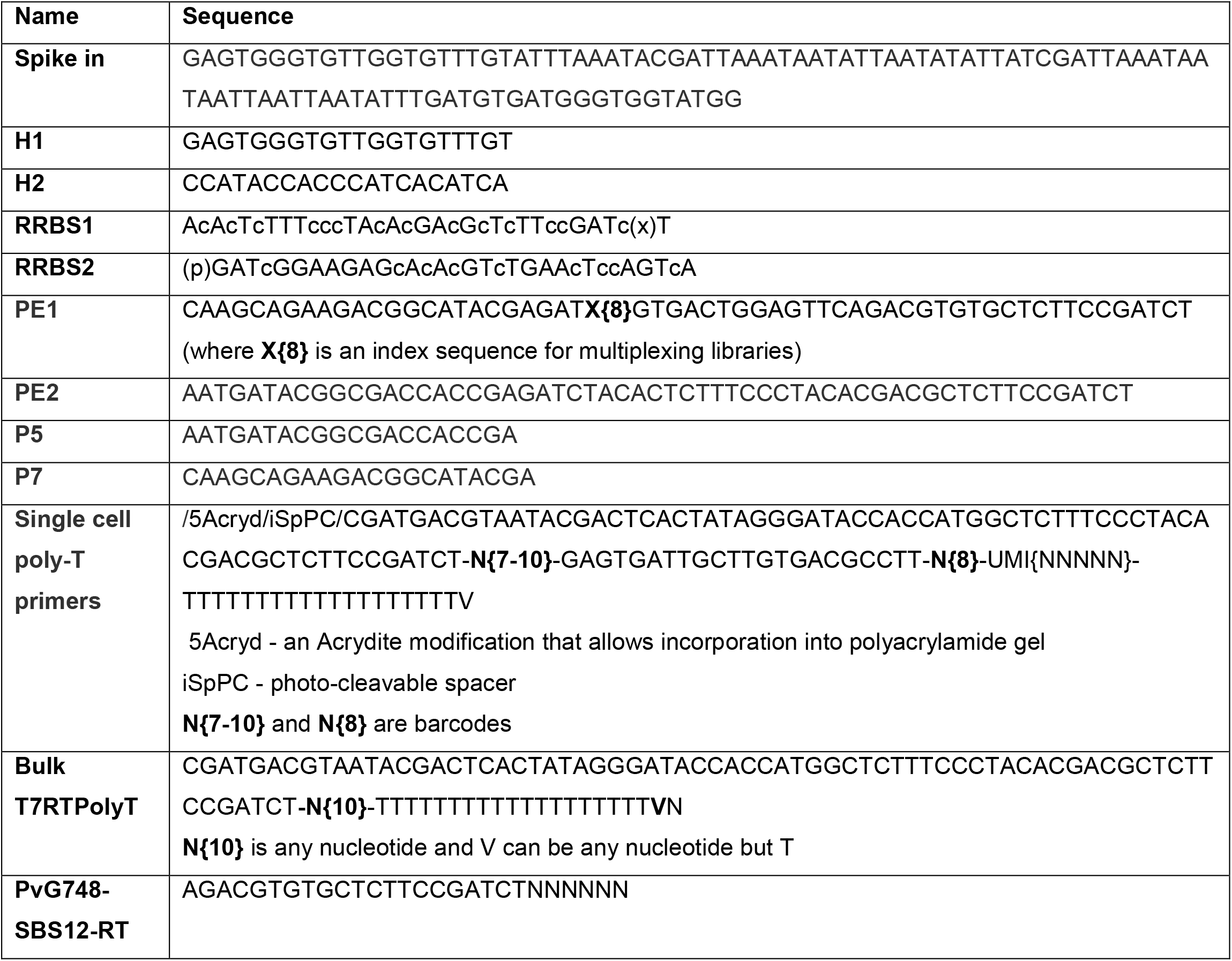
Sequences of template and primers.

For library preparation, 160 pg of spiked DNA was added to 100 ng ChIP samples to evaluate BS conversion efficiency. DNA samples were converted to blunt-ended, phosphorylated DNA using the End-It DNA End-Repair Kit (Lucigen). DNA was purified using ratio of 1:1.8 sample to AMPure XP beads. Adenosine was added to the 3’ end of the DNA fragments using Klenow (3’-5’ exo-) (New England Biolabs). DNA was purified using ratio of 1:1.8 sample to AMPure XP beads. Methylated adapters (RRBS-1, RRBS-2; Table 1) were added by ligation using DNA ligase (New England Biolabs). DNA was purified using ratio of 1:1.2 sample to AMPure XP beads. Next the DNA was subjected to bisulfite conversion using the EZ DNA Methylation Gold kit (Zymo Research) according to the manufacturer’s instructions with the addition of human total RNA as carrier. The bisulfite-converted DNA samples were amplified using KAPA HiFi HotStart Uracil (Roche) with 25 μM PE1 and PE2 full-length primers containing Illumina library indices for multiplexing (Table 1). Amplified libraries were purified using 0.7x reaction volume of AMPure XP beads and eluted in 22 μL 10 mM Tris-HCl [pH 8]. Aliquots of 15 μL of resulting libraries were run in a 2% agarose gel, and the desired 200-600 bp DNA library fragments were selected and isolated using the PureLink DNA gel extraction kit (Invitrogen). Library quality was confirmed using the Agilent 2200 TapeStation nucleic acids system and the Agilent High Sensitivity D1000 DS DNA kit. The resulting libraries had an average size of 350-550 bp. Size-selected libraries were diluted to 4 nM concentration and combined for paired-end, single index sequencing on the Illumina NextSeq 550 instrument using an illumina 550 High Output v2 (75 cycles) kit. Cycle distribution was 45 cycles for Read 1, 35 cycles for Read 2, and 8 cycles for the library index read.

### ChIP-BS-seq data analysis

Sequencing data were aligned using Bismark and Bowtie (https://www.bioinformatics.babraham.ac.uk/projects/bismark/Bismark_User_Guide.pdf) using the single-ended approach. TDF genomic browser files were produced using IGV count and data on CpGs were extracted using Bismark_methylation_extractor program, a part of the Bismark package. Using in house scripts (available upon request), we counted the number of methylated vs. non methylated CpGs within all 200-bp bins for all different chromatin marks, differentiation time points, and replicates. We applied HOMER to find peaks using ChIP-seq criteria, (we defined peak size to be 2500 bp, we turned off the option of filtering based on local and clonal signal) and used BEDTools to intersect bins with genomic intervals such as promoters, genes, and predicted enhancers.

### Single-cell experiments and library preparation

Microfluidic operations were performed using the Fluigent microfluidic pump system in a microfluidic device that was fabricated by soft lithography following standard InDrop protocols. The system contains four inlets: the cell inlet, the barcoded bead inlet, the reverse transcription/lysis mix inlet, and the carrier oil inlet. Cells were loaded into the cell inlet at 500,000 cells/ml in PBS containing 10% v/v Optiprep (Sigma) and were maintained in suspension using a magnetic micro stir bar placed in the tube. Barcoded beads were prepared as previously described ^39^ and kept in dark at 4 °C in 50% (v/v) 10 mM Tris-HCl [pH 8], 0.1 M EDTA, 0.1% (v/v)

Tween-20. For a typical experiment, 200-400 μL barcoded acrylamide beads were centrifuged in a 1.5-mL Eppendorf tube at 1000xg for 2 min to obtain a packed bead volume of 100-200 μL. After aspirating residual buffer from the pelleted beads, the tube was loaded onto the barcoded bead inlet in the microfluidics setup. The reverse transcription/lysis mix consisted of 180 μL of 5X First-Strand buffer (Invitrogen), 27 μL 10% (v/v) IGEPAL CA-630 (Sigma), 18 μL 25 mM dNTPs (NEB), 30 μL 0.1 M DTT (Invitrogen), 45 μL 1 M Tris-HCl [pH 8.0] (Biological Industries), 30 μL murine RNase inhibitor (NEB), 45 μL SuperScript III reverse transcriptase (200 U/μL, Invitrogen), and 75 μL nuclease-free water (Biological Industries) in a total volume 450 μL. The carrier oil was HFE-7500 with 3% (w/w) fluorosurfactant (RAN Biotechnologies). Typically, 2-3 mL of carrier oil were used.

During a microfluidics run, the cell suspension or encapsulated cells, reverse transcription/lysis mix, and collection tubes were kept on ice. Flow rates were 100 μL/h for cell suspension, 100 μL/h for reverse transcription/lysis mix, 25 μL/h for barcoded acrylamide beads containing photocleavable barcoding poly-T primers, and 150 μL/h for carrier oil to produce 2 nL drops. The collection tubes were exposed to 6.5 J/cm^2^ of 365-nm UV light for 10 min to release photocleavable barcoding primers from the barcoding beads. Next, the collection tubes containing the UV-exposed emulsions were transferred to a reverse transcription reaction at 50 °C for 2 h followed by 15 min at 70 °C. Each sample was demulsified by addition of 50 μL perfluoro-1-octanol (Sigma) to release the barcoded cDNA from the droplets.

To remove unused primers and primer dimers, the upper, aqueous phase containing the barcoded cDNA (~50 μL) was combined with 50 μL digestion mix containing 5 μL ExoI enzyme (NEB), 5 μL HinFI enzyme (NEB), 5 μL ExoI buffer (NEB), 5 μL CutSmart buffer (NEB), and 30 μL of nuclease-free water and incubated for 1 h at 37 °C and 10 min at 80 °C. The reaction product (in the form of a cDNA:RNA hybrid) was purified with 1.5x reaction volume of AMPure XP beads and eluted in 13.5 μL TE buffer (10 mM Tris–HCl [pH 8.0], 0.1 mM EDTA).

For second-strand synthesis, 13.5 μL digestion reaction product was combined with 1.5 μL second-strand synthesis (SSS) buffer and 1 μL of SSS enzyme mix from the NEBNext mRNA Second Strand Synthesis Module (NEB) and incubated at 16 °C for 2.5 h, followed by 20 min at 65 °C. For linear amplification by in vitro transcription, SSS reaction products (16 μL) were combined with 24 μL HiScribe T7 High Yield RNA Synthesis Kit (NEB) reagent mix containing 4 μL T7 Buffer, 4 μL ATP, 4 μL CTP, 4 μL GTP, 4 μL UTP, and 4 μL T7 enzyme mix. The reaction was incubated at 37 °C for 13 h, and the resulting RNA was purified with 1.3x reaction volume of AMPure XP beads and eluted with 20 μL TE buffer. Aliquots of 9 μL were stored at −80 °C, 2 μL were analyzed, and the remaining 9 μL were used for library preparation.

The RNA was fragmented using RNA fragmentation kit (Ambion), and 9 μL of this sample was combined with 1 μL of RNA fragmentation reagent and incubated at 70 °C for 2 min. The sample was transferred to ice, and 40 μL fragmentation stop mix containing 5 μL fragmentation stop solution and 35 μL TE buffer was added. Fragmented RNA was purified with 1.3x reaction volume of AMPure XP beads and eluted in 10 μL TE buffer. The resulting amplified and fragmented RNA was reverse transcribed using a random hexamer primer as follows: First, 10 μL RNA was mixed with 2 μL of 100 μM PvG748-SBS12-RT random hexamer primer (IDT) and 1 μL of 10 mM dNTP mix (NEB). The sample was incubated for 3 min at 65 °C and transferred to ice. The following components were added to the reaction to a total volume of 20 μL: 4 μL of 5x First-Strand SuperScript III buffer (Invitrogen), 1 μL of 0.1 M DTT (Invitrogen), 1 μL murine RNase inhibitor (NEB), and 1 μL of 200 U/μL SuperScript III reverse transcriptase (Invitrogen).

Following reverse transcription, the reaction volume was brought to 50 μL by adding 30 μL nuclease-free water, and the resulting cDNA was purified with 1.2x reaction volume of AMPure XP beads and eluted in 11.5 μL TE buffer. The resulting libraries were PCR amplified using standard PE1/PE2 full-length primer mix containing Illumina library indices for multiplexing. Each PCR reaction contained 11.5 μL post-RT cDNA library, 12.5 μL 2x KAPA HiFi HotStart PCR mix (Kapa Biosystems), and 1 μL of 25 μM PE1/PE2 index primer mix. Amplification was performed in 14 cycles. Amplified libraries were purified using 0.7x reaction volume of AMPure XP beads and eluted in 30 μL nuclease-free water. Aliquots of 15 μL of each resulting library were run in 2% agarose gels, and 200-800 bp DNA library fragments were isolated using PureLink DNA gel extraction kit (Invitrogen). Library quality was confirmed by Agilent 2200 TapeStation nucleic acids system (Agilent) using the Agilent High Sensitivity D1000 DS DNA kit. The resulting libraries had an average size of 350-550 bp. Size-selected libraries were diluted to 4 nM concentration and combined for paired-end, single-index sequencing on the Illumina NextSeq 550 instrument using an illumina 550 High Output v2 (75 cycles) kit. Cycle distribution was 45 cycles for Read 1, 35 cycles for Read 2, and 8 cycles for the library index read.

### Gene expression analysis

Single-cell RNA-seq data were processed using in-house scripts (available upon request) and analyzed using Seurat algorithm. Seurat is an R package designed for QC, analysis, and exploration of single-cell RNA-seq data (45). Cells with more than 10000 unique molecular identifiers were kept for analysis. A global-scaling normalization method “LogNormalize” with a scale factor of 10,000 was applied on the filtered dataset. Identification of highly variable genes was performed with the following parameters: x.low.cutoff =0.2; x.high.cutoff = 5; y.cutoff = 0.5 and y.high.cutoff = 10. Principal component analysis was performed on the scaled data and first 10 principal components were used for clustering and tSNE representations.

## Supplementary Figure legends

**Table S1. The number of bins (of 200 bp), peaks, and regulated genes used for analysis of ESC data.**

H3K4me1 bins are shown in green, H3K27ac bins are in red, and bins shared by both enhancer marks are in grey. Peaks were defined using the HOMER peak caller algorithm. For each row a different set of thresholds was applied; different shades mark different thresholds of 5mC percentages as indicated above each row. Genes regulated by enhancers were identified by Shen *et al.* (38).

**Table S2.** The number of bins (of 200 bp), peaks, and regulated genes used for analysis of data collected for differentiated cells.

**Table S3.** Gene list for Figure 2E.

**Table S4.** List of genes regulated by DMEs in ESCs and Diff cells.

**Table S5.** SC-RNAseq gene markers for each cluster.

**Figure S1**

Cell state validation using real-time RT-PCR (qPCR). Expression level of pluripotency markers (A) and differentiation markers (B, C) was measured. The graphs represent average gene expression levels normalized to the expression levels of GAPDH from 3 independent experiments (shown are mean values ± SEM of 3 biological replicates, asterisks represent significant changes according to 1-tailed Student’s t-test).

**Figure S2**

Boxplots of percent 5mC in ChIP-BS-seq datasets for **(A)** H3K4me1, **(B)** H3K27ac, and **(C)** H3K36me3 for ESCs and differentiated cells (Diff). The differences between the distributions of 5mC percentages are all statistically significant with p < 10^−10^ according to two-sample Wilcoxon test.

**Figure S3**

The number of LMRs with 20-80% 5mC of indicated lengths based on data from Stadler et al (23).

**Figure S4**

**(A and B)** 5mC percentages in A) ESCs (n=55,537 bins) and B) differentiated cells (Diff; n=57,093 bins) within distal enhancers bound by both H3K4me1- and H3K27ac-marked nucleosomes. The 5mC percentage is significantly higher for H3K4me1 than for H3K27ac for both cell types (t-test, p < 10^−10^). Dashed line emphasizes the differences in 5mC levels between the two marks. **(C and D)** Boxplots of 5mC percentages for each histone mark in C) ESCs and D) differentiated cells.

**Figure S5**

**(A)** 5mC percentages in H3K4me1 vs. H3K27ac in genomic segments in ESCs. Only genomic bins with more than 30 CpGs in both ChIP-BS-seq assays are shown. DMEs were defined as 200-bp bins with methylation differences exceeding 20% in either direction. **(B)** Number of H3K27ac reads for H3K4me1 DMEs and H3K27ac DMEs.

**Figure S6**

Histogram of the number of DME bins normalized by the amount of total enhancer bins in each chromosome.

**Figure S7**

Violin plots of **(A)** number of genes (nGenes) and **(B)** number of transcripts (nUMIs) calculated based on single-cell RNA-seq data using a pipeline that employs RSEM for alignment and expression profiling and the Seurat algorithm for normalization, scaling, clustering, and graphics.

**Figure S8**

Heatmap of ESCs showing most variable genes detected in Fig. 4A.

**Figure S9**

(**A**) Heatmaps showing normalized numbers of 5C and 5mC aligned reads within 200-bp bins 5 Kb up and downstream of TSSs. To reduce the number of TSSs presented, we randomly selected 3,500 TSSs with balanced representation of all expression levels. (**B**) Extraction of 5mC reads around TSSs from H3K4me1 ChIP-seq data from ESCs. Expression is plotted from high to low on the x axis for all genes (black), DME-regulated genes (green), and a randomly selected set of genes of the same size as the DME set (red). Boxplots summarizing the same data are shown in the upper right side.

**Figure S10**

MNase digestion profile analyzed by gel electrophoresis. A clear ladder, indicative of digested chromatin, is observed in both ESC replicates. Size markers in 50-bp increments are shown on the left.

