## Supplementary figures and images for "DNA Methylation Patterns Expose Variations in Enhancer-Chromatin Modifications during Embryonic Stem Cell Differentiation"

### Supplemental Figure 1

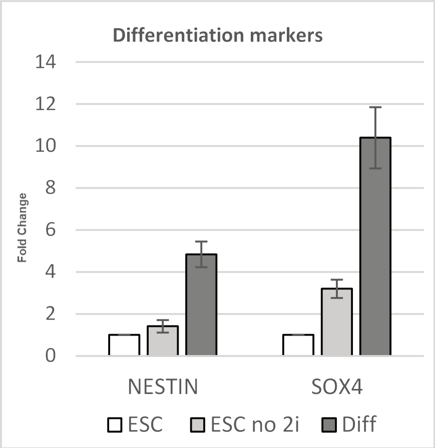

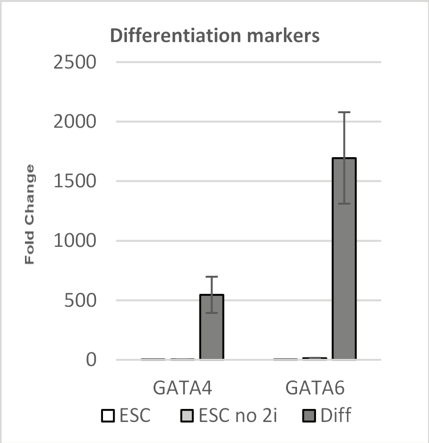

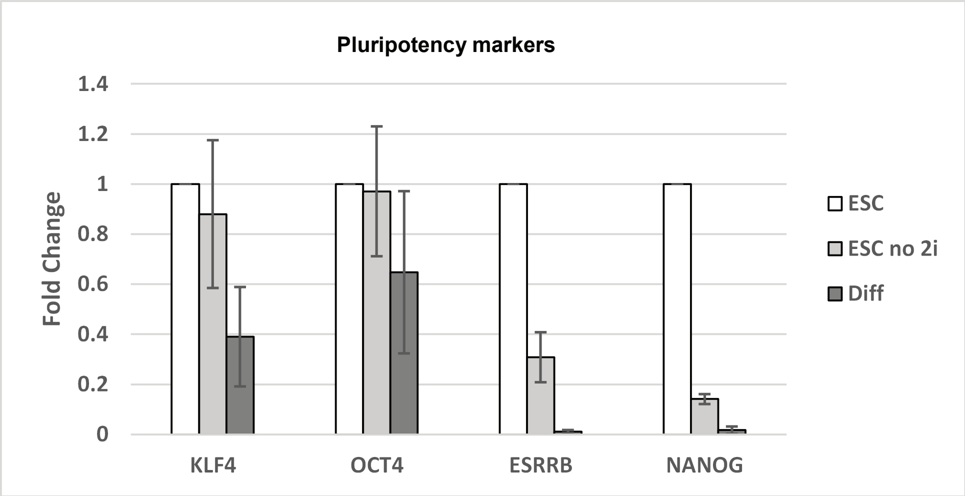

### Supplemental Figure 2

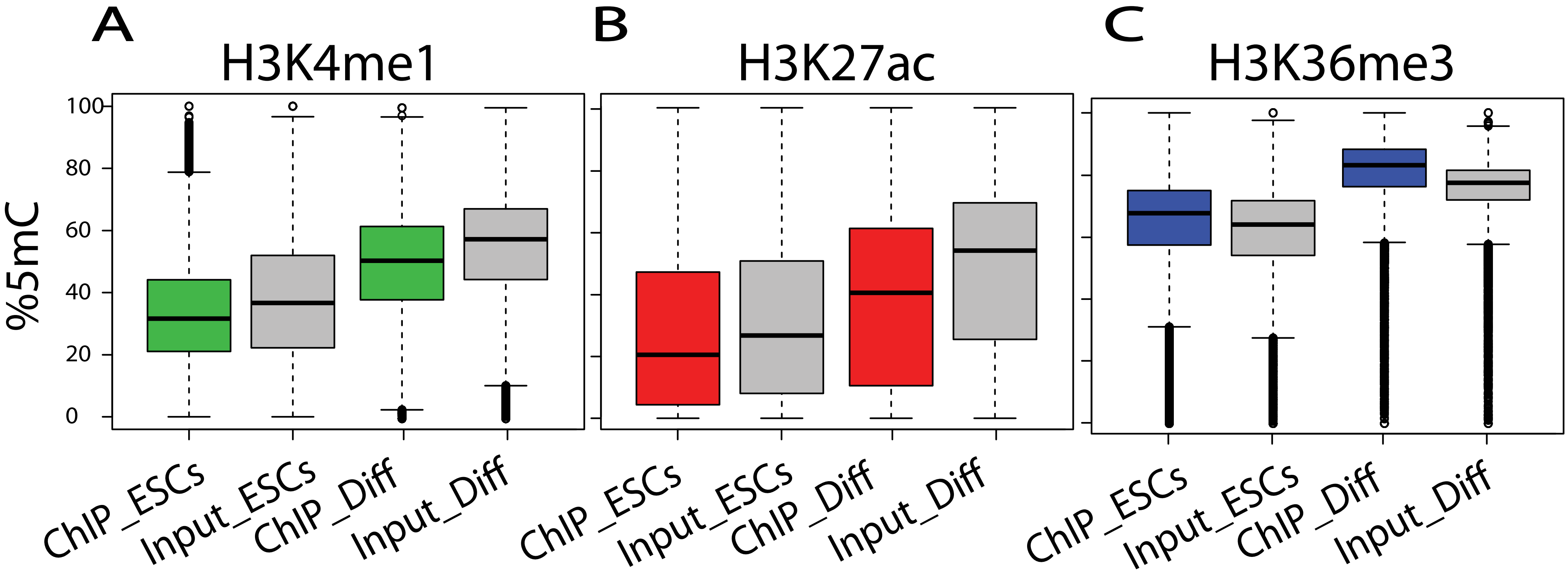

### Supplemental Figure 3

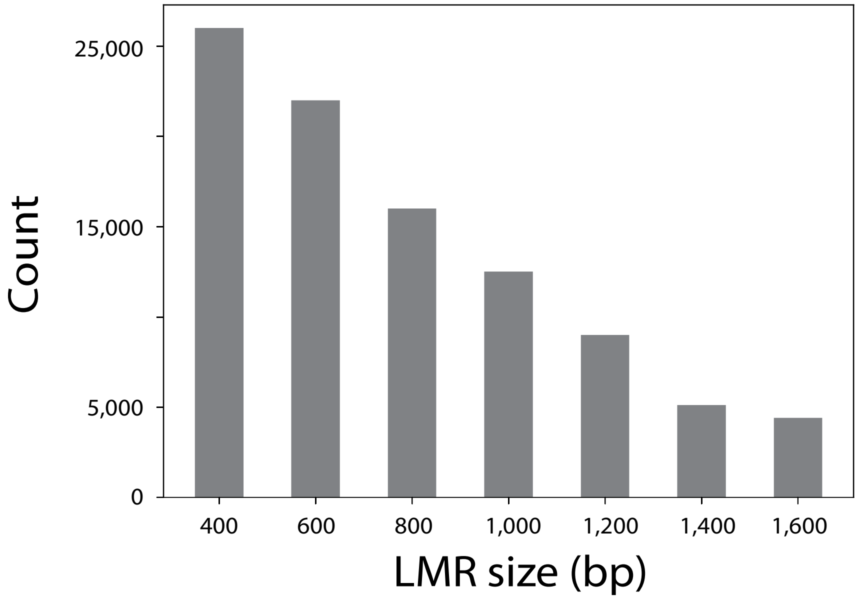

### Supplemental Figure 4

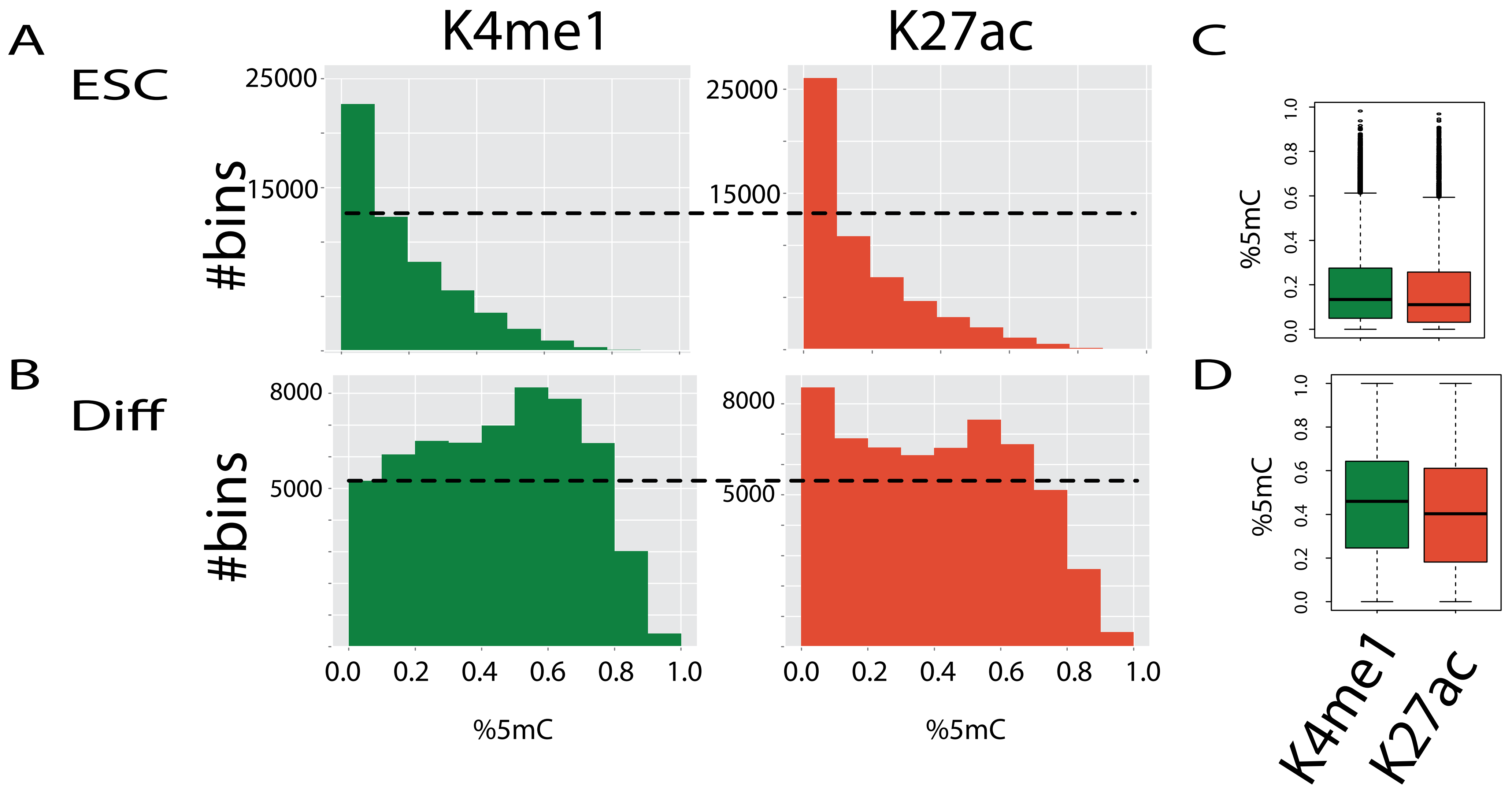

### Supplemental Figure 5

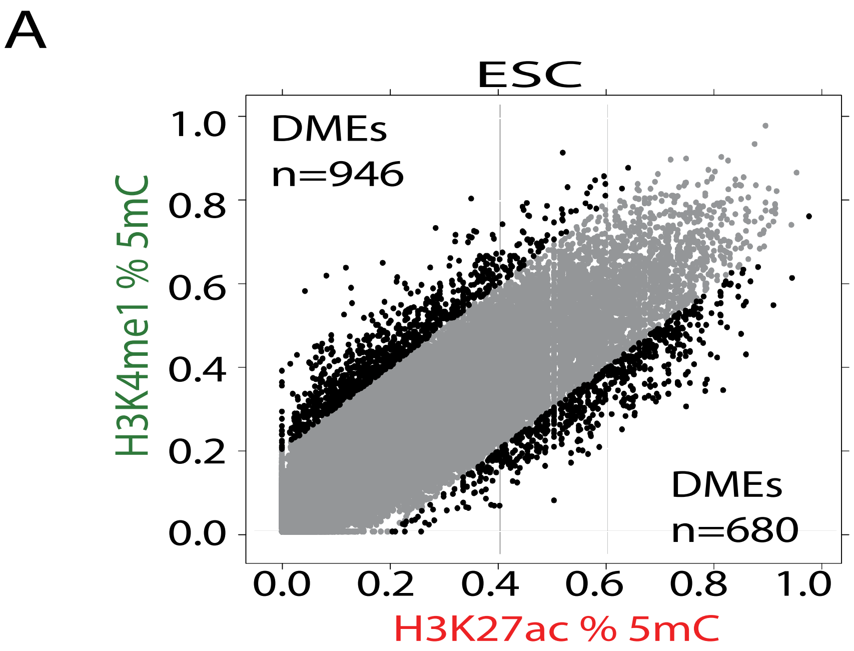

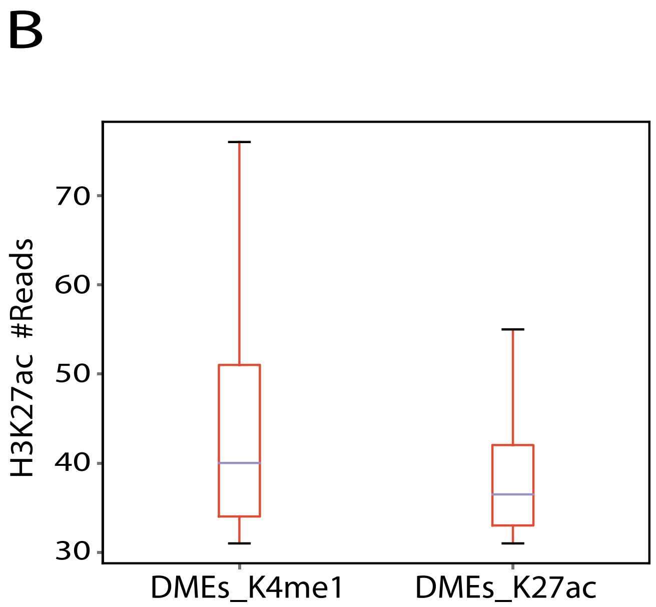

### Supplemental Figure 6

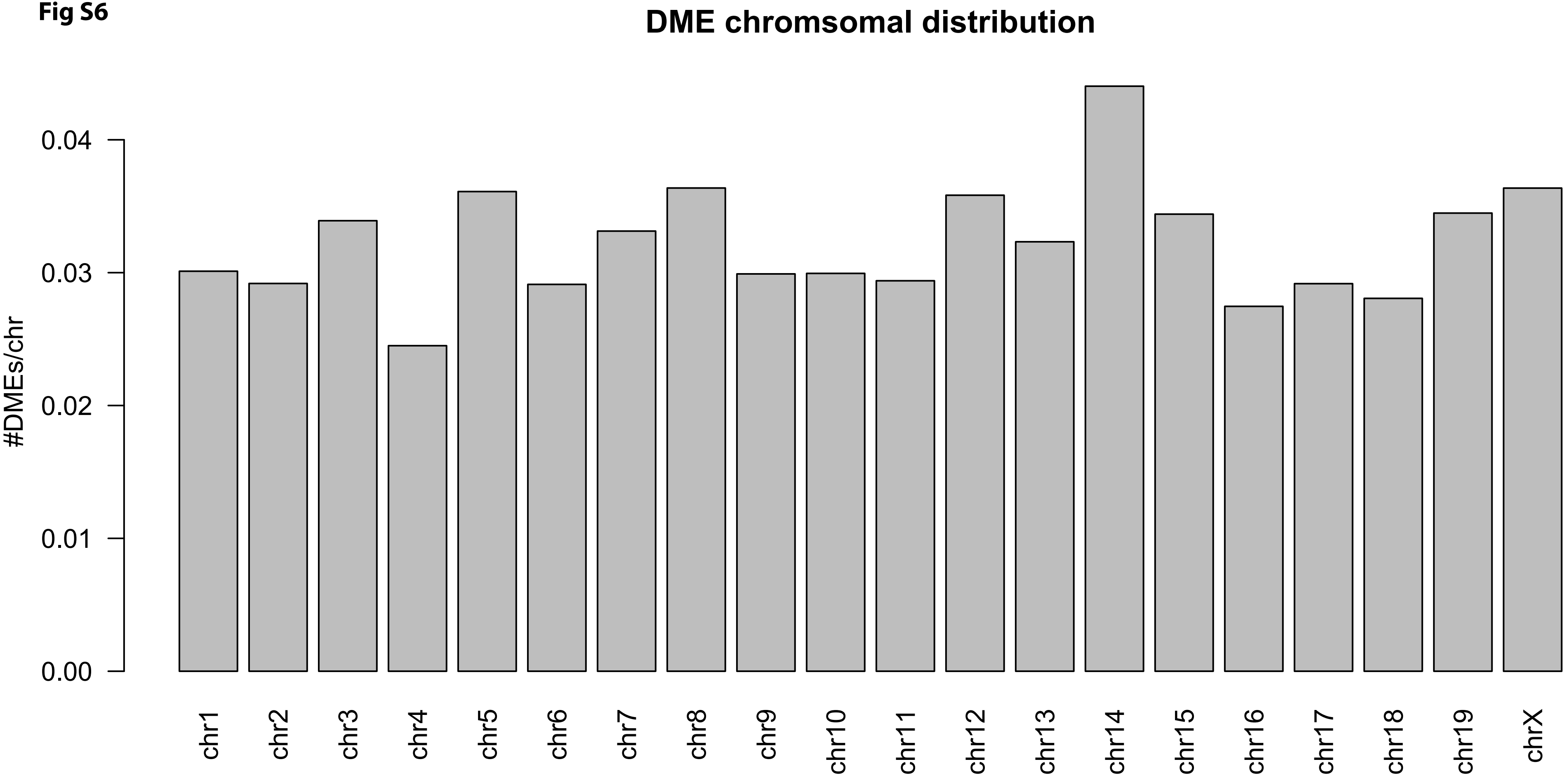

### Supplemental Figure 7

Fig S7

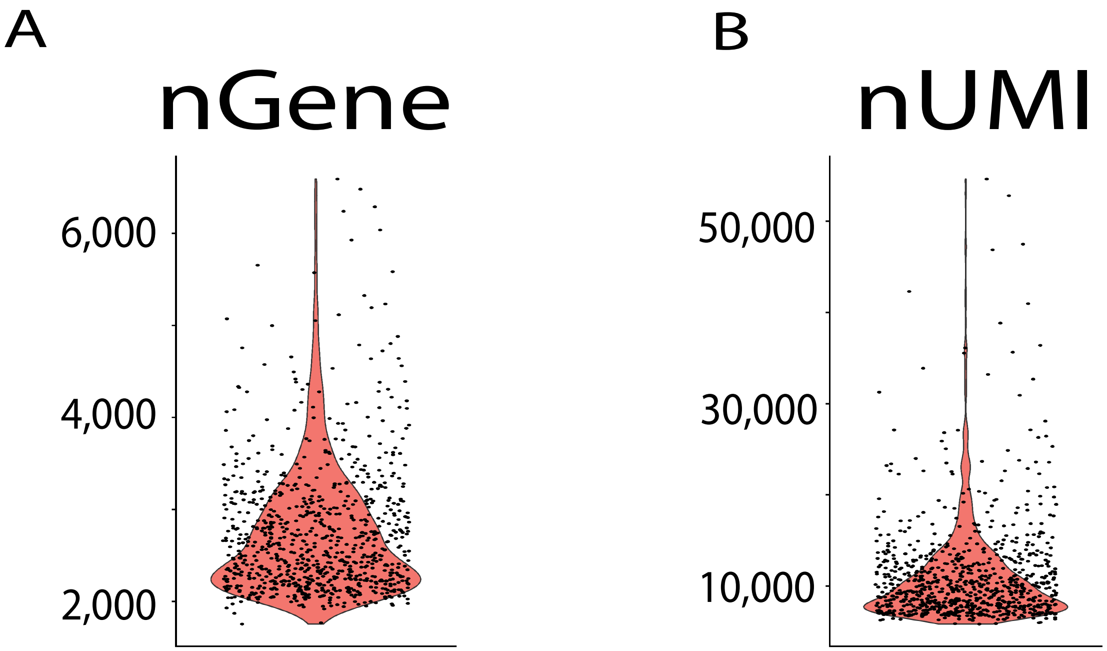

### Supplemental Figure 8

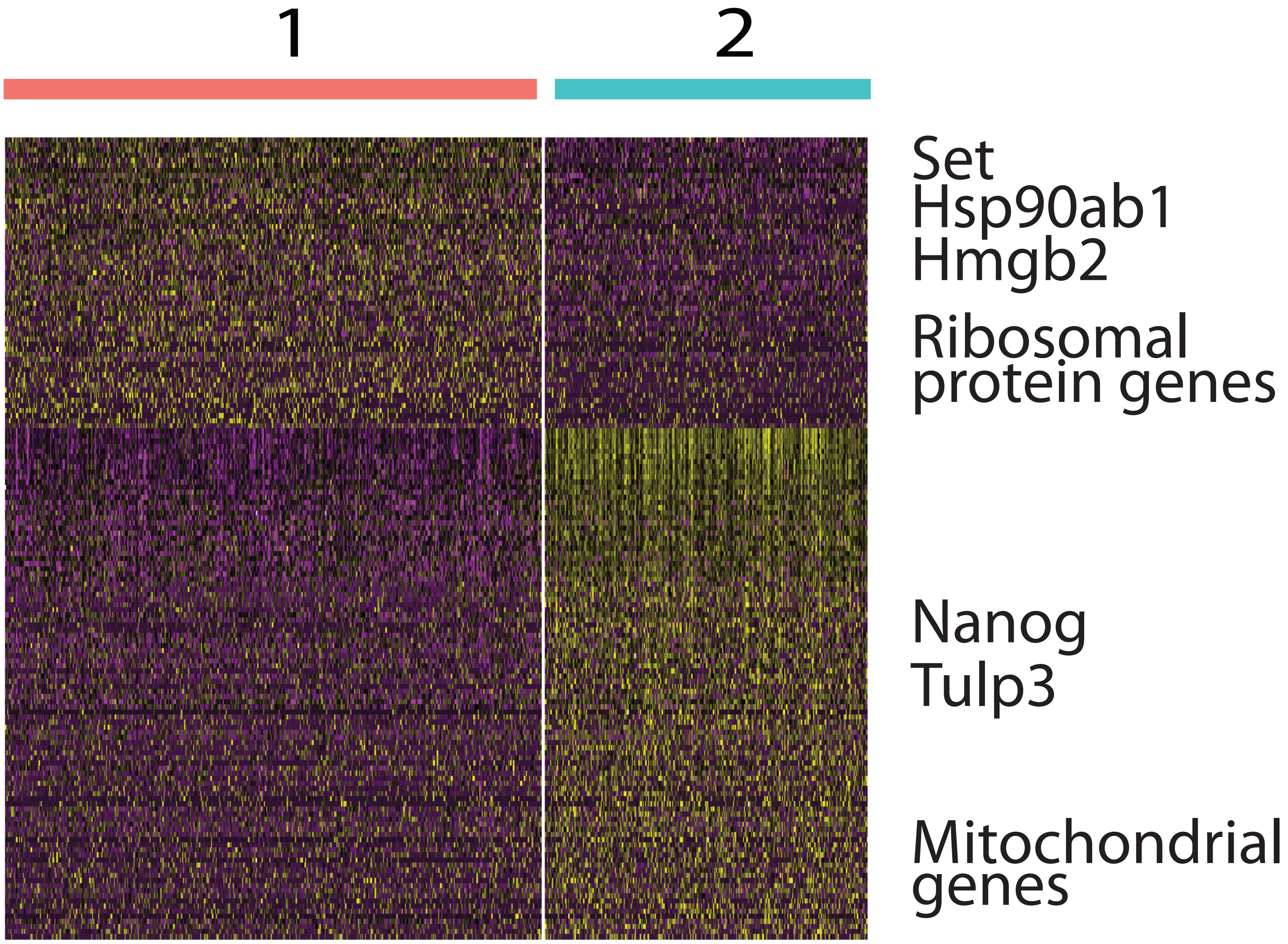

### Supplemental Figure 9

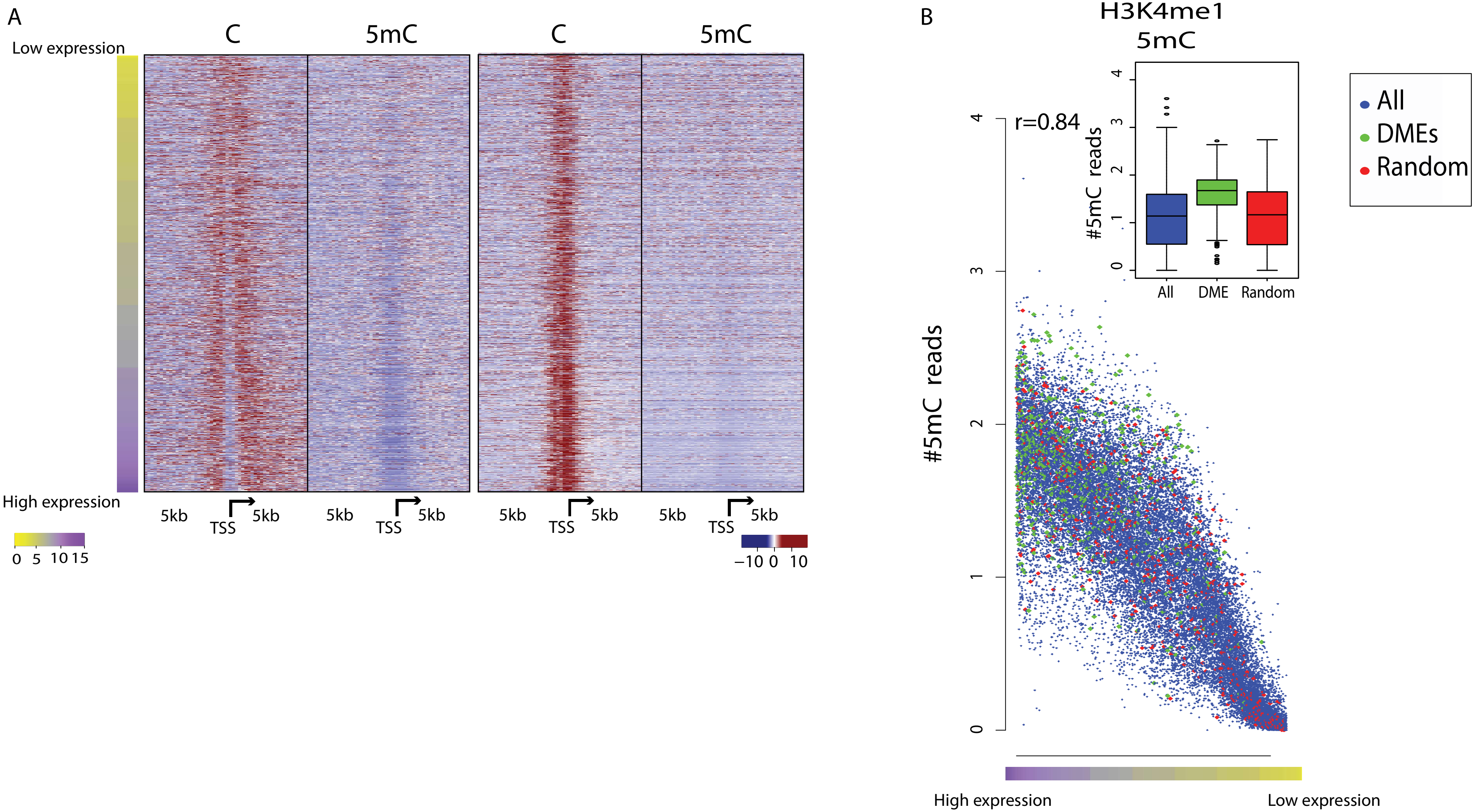

### Supplemental Figure 10

Fig S10

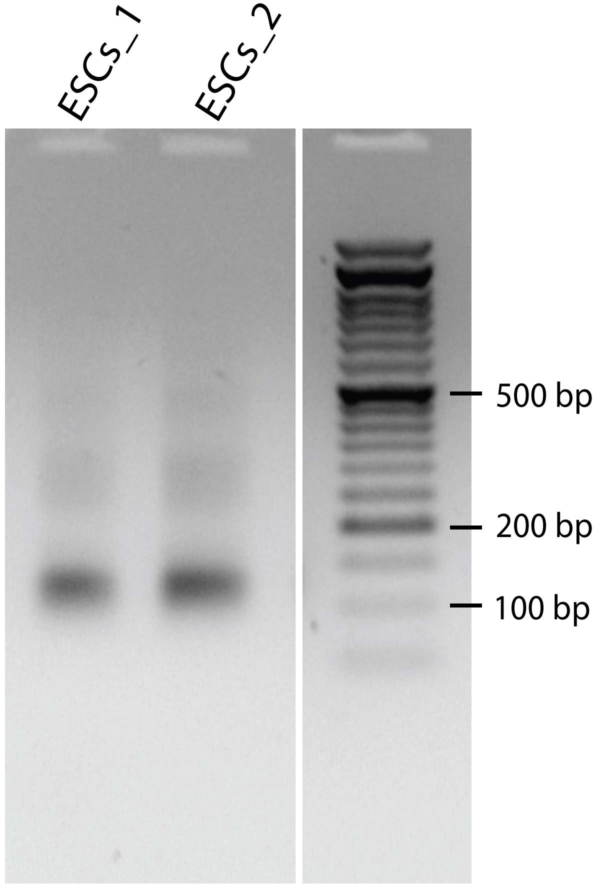
