## Supplemental Table 1 for "DNA Methylation Patterns Expose Variations in Enhancer-Chromatin Modifications during Embryonic Stem Cell Differentiation"

**Table S1**

### ChIP-BS-Seq data from ESCs

## H3K4me1

| | Bins | Peaks | Regulated Genes |
|---|---|---|---|
| | 1,308,520 | 51,582 | 11,976 |
| Bins CpG>30 | 356,366 | 44,946 | 11,299 |
| Bins CpG>30 %5mC<20 | 107,845 | 24,868 | 5,260 |
| Bins CpG>30 %5mC>30 | 28,462 | 12,701 | 5,132 |

## H3K27ac

| | Bins | Peaks | Regulated Genes |
|---|---|---|---|
| | 1,230,054 | 36,661 | 9,248 |
| Bins CpG>30 | 66,210 | 16,356 | 6,170 |
| Bins CpG>30 %5mC<20 | 45,340 | 10,848 | 4,850 |
| Bins CpG>30 %5mC>30 | 4,932 | 3,736 | 1,963 |

#### H3K4me1 and H3K27ac

| | Bins | Peaks | Regulated Genes |
|---|---|---|---|
| | 1,158,451 | 23,488<br>28,913 | 8,209 |
| Bins CpG>30 | 55,537 | 12,025<br>13,226 | 5,493 |
