## Supplemental Table 2 for "DNA Methylation Patterns Expose Variations in Enhancer-Chromatin Modifications during Embryonic Stem Cell Differentiation"

Table S2

### ChIP-BS-Seq data from differentiated cells

## H3K4me1

| | Bins | Peaks | Regulated Genes |
|---|---|---|---|
| | 1,270,711 | 43,615 | 10,733 |
| Bins CpG>30 | 257,891 | 37,918 | 9,927 |
| Bins CpG>30 %5mC<20 | 25,092 | 10,513 | 2,939 |
| Bins CpG>30 %5mC>30 | 76,423 | 21,729 | 7,146 |

## H3K27ac

| | Bins | Peaks | Regulated Genes |
|---|---|---|---|
| | 1,204,246 | 27,858 | 7,957 |
| Bins CpG>30 | 64,080 | 15,904 | 5,929 |
| Bins CpG>30 %5mC<20 | 19,689 | 6,713 | 3,484 |
| Bins CpG>30 %5mC>30 | 13,830 | 7,293 | 3,536 |

#### H3K4me1 and H3K27ac

| | Bins | Peaks | Regulated Genes |
|---|---|---|---|
| | 1,108,529 | 19,497<br>23,889 | 7,239 |
| Bins CpG>30 | 57,093 | 12,287<br>14,122 | 5,375 |
